# Optimized ChIP-exo for mammalian cells and patterned sequencing flow cells

**DOI:** 10.1101/2025.08.14.670389

**Authors:** Daniela Q. James, Sohini Mukherjee, C. Caiden Cannon, Shaun Mahony

## Abstract

By combining chromatin immunoprecipitation (ChIP) with an exonuclease digestion of protein-bound DNA fragments, ChIP-exo characterizes genome-wide protein-DNA interactions at near base-pair resolution. However, the widespread adoption of ChIP-exo has been hindered by several technical challenges, including lengthy protocols, the need for multiple custom reactions, and incompatibilities with recent Illumina sequencing platforms. To address these barriers, we systematically optimized and adapted the ChIP-exo library construction protocol for the unique requirements of mammalian cells and current sequencing technologies. We introduce a Mammalian-Optimized ChIP-exo (MO-ChIP-exo) protocol that builds upon previous ChIP-exo protocols with systematic optimization of crosslinking, harvesting, and library construction. We validate MO-ChIP-exo by comparing it to previously published ChIP-exo protocols and demonstrate its adaptability to both suspension (K562) and adherent (HepG2, mESC) cell lines. This improved protocol provides a more robust and efficient method for generating high-quality ChIP-exo libraries from mammalian cells.

**SUMMARY:** ChIP-exo is a genome-wide protein-DNA binding assay with unrivalled resolution, but its widespread adoption has been hindered by technical challenges, particularly when applied to mammalian cells or when used with recent sequencing platforms. We introduce a Mammalian-Optimized ChIP-exo (MO-ChIP-exo) protocol with key modifications that overcome previous technical hurdles. We demonstrate that our optimized protocol produces high-quality data comparable to previously published protocols and is adaptable for use with both suspension (K562) and adherent (HepG2, mESC) cell lines.

## INTRODUCTION

Carefully orchestrated protein-DNA interactions mediate critical cellular processes, including gene regulation, genome organization, replication, and DNA repair. Characterizing the genome-wide binding locations of transcription factors and other regulatory proteins is thus crucial for understanding these complex cellular functions. Chromatin immunoprecipitation (ChIP) greatly advanced the study of protein-DNA interactions by enabling the identification of specific binding events *in vivo* (Gilmour and Lis 1984). ChIP involves cross-linking proteins to DNA, followed by the isolation of specific protein-DNA complexes using an antibody against a protein of interest. This selective immunoprecipitation allows for examination of the DNA sequences associated with the targeted protein. ChIP combined with high throughput sequencing (ChIP-seq) (Johnson et al. 2007; Albert et al. 2007) emerged as the main method to identify genome-wide patterns of protein-DNA binding. ChIP-seq has been deployed in a wide variety of species and cell types to map the localization of histone modifications and variants, chromatin remodelers, and transcription factors (Feingold et al. 2004; Moore et al. 2020). While newer approaches such as Cleavage Under Targets & Release Using Nuclease (CUT&RUN) (Skene and Henikoff 2018) and Cleavage Under Targets and Tagmentation (CUT&Tag) (Kaya-Okur et al. 2019) have provided valuable alternatives, they are most popularly applied to characterize histone modifications (Abbasova et al. 2025). ChIP-seq remains the preferred method to study mammalian transcription factor binding.

ChIP-exo (Rhee and Pugh 2011; 2012) greatly improved upon the resolution of ChIP-seq by incorporating an exonuclease to digest immunoprecipitated DNA fragments in a strand-specific 5′ → 3′ direction. The exonuclease proceeds until blocked by a crosslinked protein, or fully digests fragments not crosslinked to a protein. As a result, ChIP-exo yields sharper protein-bound fragments and lower background than ChIP-seq, enabling the identification of binding events at near base-pair resolution. ChIP-exo was applied to study transcription factor binding in yeast (Rhee and Pugh 2011) and to characterize the organization of individual histones in the yeast genome (Rhee et al. 2014). Most notably, ChIP-exo was used to characterize the genome-wide architecture of all yeast chromatin-associated proteins in the Yeast Epigenome Project (Rossi et al. 2021). It has also been applied to screen a large cohort of monoclonal antibodies generated by the Protein Capture Reagents Program (PCRP) (Lai et al. 2021).

Despite its advantages, the widespread adoption of ChIP-exo has been hindered by several technical challenges, including lengthy protocols, the need for multiple custom reactions, a lack of quality control checkpoints, and higher costs. Previous efforts have attempted to address these issues. The original protocol was initially adapted from the SOLiD (ChIP-exo 1.0) to the Illumina sequencing platform (ChIP-exo 1.1) (Serandour et al. 2013; Yen et al. 2013). The ChIP-nexus protocol adapted ChIP-exo to incorporate more efficient adapter ligation and library amplification strategies with single-end sequencing (He et al. 2015). Most recently, Rossi, et al., presented a simplified protocol (ChIP-exo 5.0), which greatly reduced the number of enzymatic steps while producing data comparable to earlier methods (Rossi et al. 2018). However, these previous efforts were tailored to the technologies available at the time, including random patterned flow cells, combinatorial index IDs for multiplexing, and gel excisions for size selection. The advent of newer Illumina platforms, such as the NextSeq 2000, which use patterned flow cells and exclusion amplification clustering, has introduced new complications. These technologies have revealed and potentially exacerbated issues in the ChIP-exo 5.0 protocol, including index hopping and a high occurrence of poly-G sequences in paired-end sequencing libraries (where the latter results from the inability of the Illumina two-color chemistry to distinguish between a string of ‘G’ bases and ‘no signal’). Furthermore, the earlier protocols were primarily optimized to target yeast or *Drosophila* and have not been systematically optimized for mammalian cells.

These challenges have prompted us to re-evaluate and adapt the ChIP-exo library construction protocol, specifically considering recent advancements in sequencing technology and the unique requirements of mammalian cells. In this manuscript, we highlight key protocol modifications for generating ChIP-exo libraries from mammalian cell lines. We present a series of improvements developed through the systematic exploration of multiple conditions for cell harvesting and library construction, building upon the foundation of the ChIP-exo 5.0 protocol.

## RESULTS

### Overview of the ChIP-exo protocol

The ChIP-exo 5.0 protocol offered a simplified series of steps compared with the original ChIP-exo protocol (Rossi et al. 2018) (**Figure 1**). ChIP-exo 5.0 begins with crosslinking, where cells are treated with 1% formaldehyde in PBS for 10 minutes at RT. The crosslinking reaction is quenched with a 125 mM Glycine solution for 5 minutes. After washing in PBS, pellets are snap-frozen in liquid nitrogen. Crosslinked cells undergo sequential cytoplasmic and nuclear lysis in the presence of a complete protease inhibitor before chromatin is sheared via sonication in IP buffer (containing SDS) for 10 cycles to obtain DNA fragments 100 to 500 bp in size. Before proceeding to the immunoprecipitation (IP) step, the sheared chromatin (10 M cell equivalent) is first pre-cleared to reduce background noise. The IP reaction proceeds by incubating chromatin with magnetic beads that are covalently linked to Protein A or G and pre-conjugated to an antibody targeting the protein of interest. Following stringent washes, bead-bound chromatin is subjected to A-tailing on the 3′ ends in preparation for the first adapter ligation (in ChIP-exo 1.1 a polishing step preceded the A-tailing but this was removed in ChIP-exo 5.0). T4 DNA ligase is used to add the Read 2 sequencing adapter. After this adapter has been ligated, a fill-in reaction fills in the 5′ overhang. The defining step of the protocol, lambda exonuclease digestion, is then performed. This enzyme digests single strands of DNA in the 5′ → 3′ direction until it is blocked by a crosslinked protein. Next, crosslinks are reversed, the sample is treated with proteinase K, and DNA is purified. The Read 1 sequencing adapters are attached to the 5′ ends of the digested single-stranded molecules via splint ligation. Because the Read 1 sequencing adapters are added to the side of the molecule that was digested with exonuclease, Read 1 sequencing will generate the precise DNA footprints that are characteristic of ChIP-exo’s high resolution. Read 2 is the product of the first adapter ligation following A-tailing, making it equivalent to a ChIP-seq experiment in resolution. Finally, fragments are enriched through PCR (18 cycles). Libraries are assessed by amplifying a quarter of the reaction for an additional six cycles (24 total) and confirming their presence by electrophoresis. Completed libraries (18 cycles) are size selected by performing a gel excision of fragments in the 200 to 500 bp region followed by DNA cleanup using a gel extraction kit.

**Figure 1:**
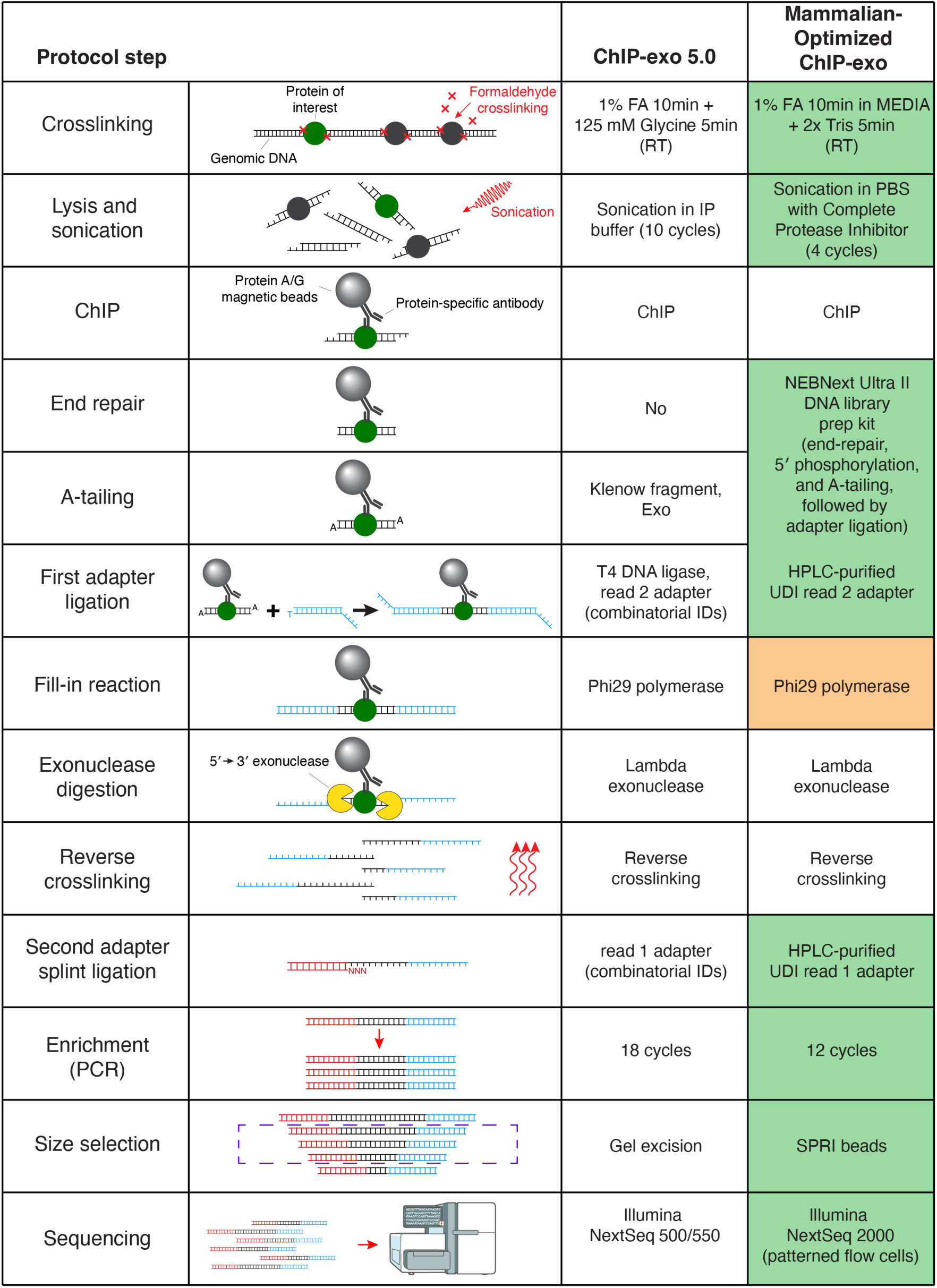
Summary of the ChIP-exo protocol, highlighting MO-ChIP-exo protocol modifications. Each row describes a key step in the ChIP-exo protocol. The second last column describes protocol steps in ChIP-exo 5.0, while the last column describes steps in MO-ChIP-exo. The shaded boxes in the last column highlight key protocol steps that were modified (green) or explored (orange) in MO-ChIP-exo.

Our initial attempts to execute the above ChIP-exo 5.0 protocol on mammalian cells identified several inefficiencies and some incompatibilities with newer patterned flow cell sequencing platforms. We therefore set out to reevaluate various steps in the protocol with the goal of optimizing for mammalian cells and assessing the efficiencies of techniques that were unavailable when ChIP-exo 5.0 was defined. The highlighted cells in **Figure 1** summarize the protocol steps that we re-assessed in this study, resulting in the mammalian-optimized ChIP-exo protocol (MO-ChIP-exo). The following sections will describe each of these optimizations in turn.

Our investigations primarily focus on ChIP-exo experiments targeting the CCCTC-binding factor (CTCF) in K562. CTCF is a highly conserved zinc finger protein best known as an insulator transcription factor that helps to establish chromosome architecture (Kim et al. 2015). CTCF offers a convenient target for quality assessments, given the availability of high-quality antibodies, its ubiquitous expression in mammalian cells, and the high conservation of CTCF binding sites across cell types (H. Chen et al. 2012). We use a series of quality metrics to compare sequenced ChIP-exo library quality across multiple conditions. **<u>F</u>raction of <u>R</u>eads in <u>P</u>eaks (FRiP)**, defined by ENCODE as the fraction of mapped reads that overlap peak regions, was used to assess ChIP enrichment and signal-to-noise (Landt et al. 2012). To facilitate comparisons of FRiP scores across samples, we measure it in relation to a common set of ENCODE CTCF peaks (see **Methods**). **<u>E</u>stimated <u>L</u>ibrary <u>S</u>ize (ELS)** is an empirical estimate of the number of unique DNA fragments present in each sequencing library, thus serving as a library complexity measurement. Finally, the **Peak Counts** for each sample are reported by the ChExMix peak-finder for ChIP-exo data (Yamada et al. 2018; 2020), and represent the numbers of genomic regions that display a statistical enrichment of aligned reads in the ChIP-exo sample compared with a control.

### Optimizing crosslinking conditions for high-quality ChIP-exo

*In vivo* crosslinking, the first step in the ChIP-exo protocol, is critical for preserving protein-DNA interactions in a stable yet reversible manner. This reversibility is critical for isolating specific chromatin complexes of interest and later analyzing their components. While crosslinking is a pivotal step for ChIP, the ideal conditions have been debated for decades (reviewed in Sutherland et al. 2008; Hoffman et al. 2015). Optimal crosslinking balances the resolution of binding site detection with epitope loss from shearing. Because crosslinking efficiency is affected by the crosslinking agent as well as reaction pH, temperature, time, and concentration (Baranello et al. 2016; Ji et al. 2022; Xu et al. 2024), these conditions must be fine-tuned to detect biologically relevant interactions. Too little crosslinking will only capture the most robust interactions (Kasinathan et al. 2014), while too much crosslinking can fix even the most transient interactions, leading to insoluble complexes that mask or reduce epitope accessibility (Rozbeský et al. 2018). Formaldehyde is the most widely used crosslinking agent due to its cell permeability, relatively fast reactivity, low cost, and ability to crosslink molecules in close proximity. It diffuses through the cell membrane and forms covalent bonds between nucleophilic groups on proteins and DNA. Despite its widespread use, protocols vary in terms of the crosslinking vehicle (which may influence pH stability), temperature, formaldehyde concentration, and exposure time. The pH is a critical variable, as it affects both crosslinking reactions (Schroeder et al. 2019) and protein affinity (Akerström and Björck 1986; Watanabe et al. 2009). Buffers like PBS are commonly used to sustain a physiological pH (Keller et al. 2021; Park et al. 2025), but many studies either crosslink directly in cell growth media (Giresi et al. 2007) or fail to specify the vehicle used (Euskirchen et al. 2007; Johnson et al. 2007; Quon et al. 2023). Although practical from a technical standpoint, crosslinking cells in media introduces variations from the media components, which can alter pH or quench formaldehyde.

Our initial attempts to produce ChIP-exo libraries using anti-CTCF antibody by crosslinking cells in PBS at room temperature resulted in poor quality metrics, including low FRiP scores, low library complexity, and low numbers of peaks (**Figure 2A, 2B**). We directly compared these libraries to libraries from cells crosslinked in rich media (IMDM+10%FBS) and observed a striking improvement, with higher FRiP scores, larger ELS, and a greatly increased number of peaks. The improvement observed from crosslinking cells in media as compared to PBS was recapitulated independently with an anti-USF1 antibody (**Supplementary Figure 1**).

**Figure 2.**
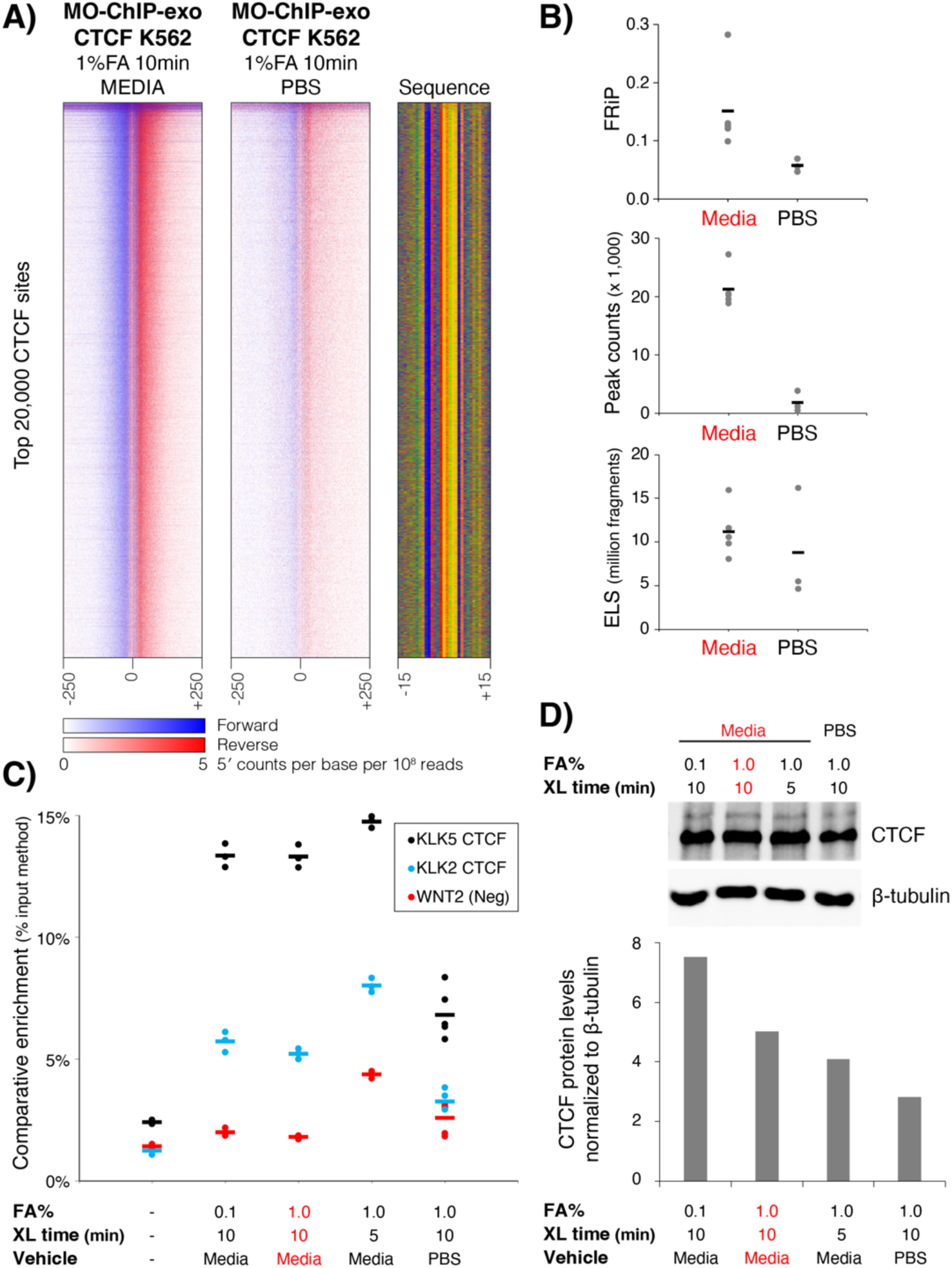
Crosslinking vehicle influences reaction efficiency and CTCF library quality. **A)** Heatmaps display normalized per base enrichment of 5′ read positions from ChIP-exo libraries constructed from cells crosslinked in media (left) and PBS (right). The heatmaps plot enrichment in 500bp windows centered on the top 20,000 CTCF peaks. Blue represents forward strand 5′ enrichment, while red represents reverse strand 5′ enrichment. **B)** Library quality metrics from ChIP-exo libraries constructed from cells crosslinked in media and PBS, including FRiP scores, peak counts, and estimated library sizes. Each data point represents an individual ChIP-exo library and short lines represent the mean value across replicates. **C)** ChIP-qPCR was used to quantify comparative enrichment of CTCF at binding sites in the KLK locus (KLK2 and KLK5) and in an unbound negative control region (WNT2) across multiple crosslinking conditions. Dots represent individual technical replicates (minimum 3 per condition) and short lines represent mean values across replicates. **D)** Western blot (top panel) showing CTCF protein levels for K562 cells across multiple crosslinking conditions. And quantification of CTCF protein levels normalized to Beta-tubulin (bottom panel) from the Western blot. MO-ChIP-exo conditions are highlighted throughout in red.

To understand the differences observed between cells crosslinked in media and PBS, we examined the quality of the input chromatin. First, we compared the DNA yield for each condition after reverse crosslinking sonicated chromatin (i.e., before ChIP), finding a higher yield when cells were crosslinked in media (**Supplementary Figure 2A**). This difference can be partly explained by the elimination of a centrifugation step needed to resuspend the cells in PBS, which affects yield by almost 50% without affecting viability (**Supplementary Figure 2D-2F**). Interestingly, the fraction of DNA in the target fragment size range (100-500bp) did not differ substantially between media and PBS conditions (**Supplementary Figure 2B**). Next, we examined the CTCF-ChIPed material and determined that cells crosslinked in media yielded a significantly higher recovery of CTCF-bound DNA (**Supplementary Figure 2C**). We also used ChIP-qPCR to quantify CTCF binding at two constitutive binding sites near the KLK2 and KLK5 gene loci (Khoury et al. 2020), compared to a locus at the WNT2 gene with no known CTCF binding. CTCF enrichment is significantly higher in the material collected from cells crosslinked in media compared to cells crosslinked in PBS (**Figure 2C**) (one-tailed T-test, KLK2 *p* = 4.6×10^-5^, KLK5 *p* = 1.12×10^-6^). This enrichment was maintained even when formaldehyde concentration was decreased tenfold or crosslinking time was halved (5 min instead of 10 min) (**Figure 2C**). Finally, we used Western blots to assess the recovery of CTCF protein from K562 whole cell lysates after crosslinking, finding that the amount of CTCF detected was higher for the samples crosslinked in media (**Figure 2D**). Together, these results show that the crosslinking vehicle affects chromatin yield and potentially affects integrity of DNA-protein interactions or epitope accessibility, with crosslinking in media leading to improved ChIP quality and reduced background noise.

After establishing media as the optimal crosslinking vehicle, we assessed other reaction variables. We found that a formaldehyde concentration of 1% yielded the best library quality (**Figure 3A**), whereas formaldehyde concentrations of 0.1% or 0.83% produced libraries with lower FRiP scores and low numbers of peaks (**Figure 3A**). Reducing the crosslinking time from 10 to 5 minutes did not have a large effect (**Figure 3A**). We also compared the quenching agents Tris and Glycine, which are added in excess to halt the crosslinking reaction. While Glycine is typically used to quench formaldehyde, Tris is a more efficient quencher given each molecule’s ability to react with 2 formaldehyde molecules (reviewed in (Hoffman et al. 2015)). However, both quenchers produced libraries of comparably high quality (**Figure 3A**). We noted that both quenchers, but particularly Glycine, lowered the media’s pH. Specifically, the initial pH of the media (pH 6.9 to 7.3) was unchanged after adding formaldehyde but decreased to 5.2 to 6.3 after the addition of powder or liquid Glycine and to 6.5 to 6.7 after the addition of Tris (**Figure 3B**). This pH drop may contribute to Glycine’s effectiveness by slowing the reaction rate. Despite their similar performance, we chose Tris for the MO-ChIP-exo protocol due to its greater theoretical quenching capability, which minimizes the risk of over-crosslinking. Finally, we confirmed that quenching conditions were unable to mitigate the detrimental effects of crosslinking in PBS (**Supplementary Figure 3**).

**Figure 3.**
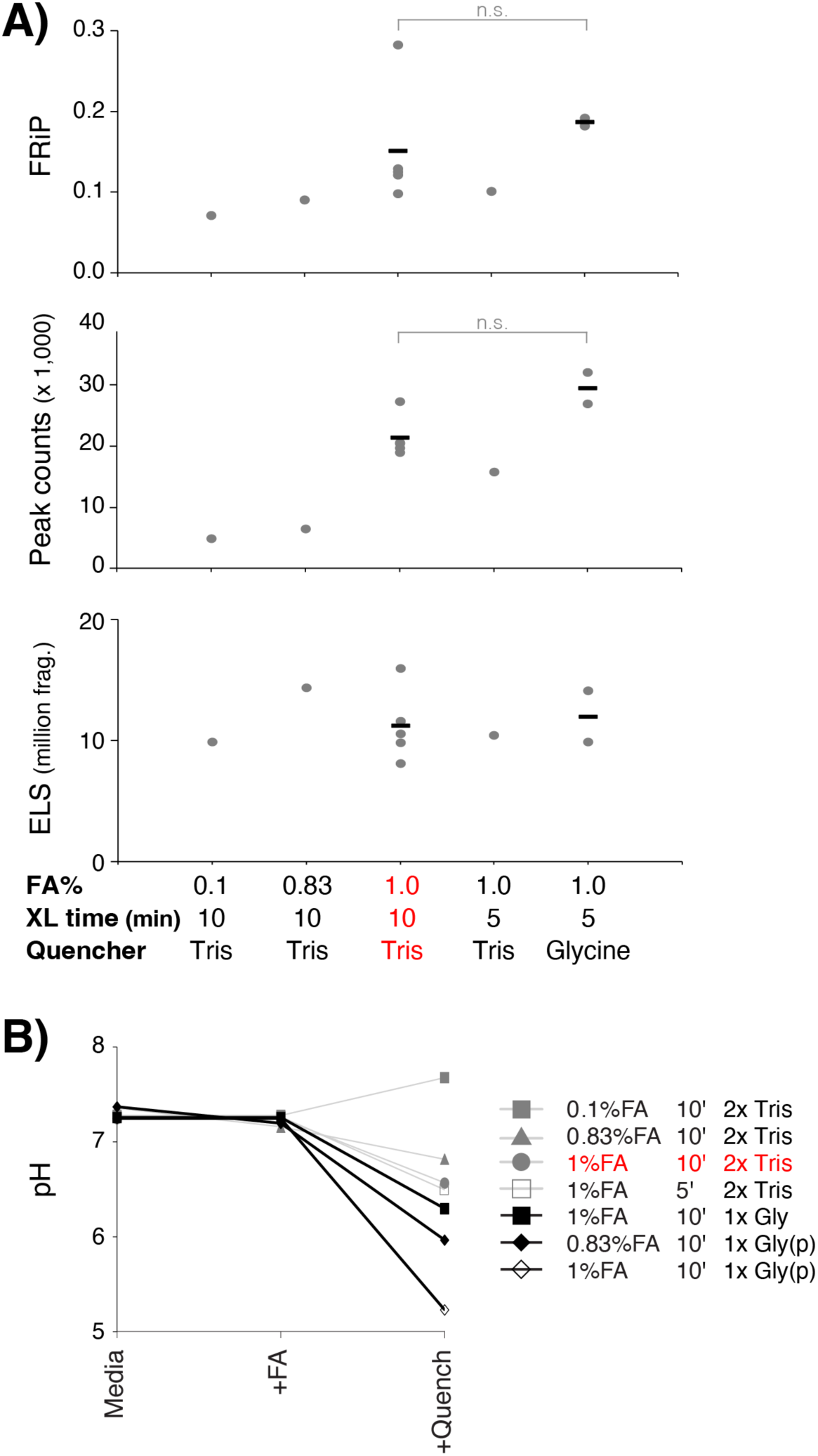
Formaldehyde concentration, time, and quenchers influence ChIP-exo library quality. **A)** Quality metrics are shown for CTCF libraries made under different crosslinking conditions in media (0.1%, 0.83%, or 1% FA for 10 min or 1% FA for 5min) using Tris or Glycine as quenchers. There is no significant difference between the indicated Tris and Glycine conditions (T test; FRiP *p* = 0.17; peak counts *p* = 0.14).**B)** pH for media before and after crosslinking and quenching using multiple FA concentrations and times as well as different quenchers. MO-ChIP-exo conditions are highlighted throughout in red.

Crosslinking immobilizes cell scaffold/topology, which eventually affects cell function and adhesion and inevitably leads to cell death. Fast crosslinking is expected to preserve interactions without immediate effects to viability, making crosslinking time a critical factor. Trypan blue staining confirmed that crosslinking as performed in MO-ChIP-exo [(10min, 1% formaldehyde), quenching (5min, Tris) and washing (cold PBS)] does not immediately affect cell viability (**Supplementary Figure 2F**) despite observations of adherent cells becoming dislodged in the presence of 1% formaldehyde with gentle shaking. Further, to preserve chromatin integrity after quenching and to minimize the possibility of proteolysis, we introduce protease inhibitors during harvest and we snap freeze the cell pellets using liquid nitrogen. The liquid nitrogen snap freeze is conducive to cell lysis which is performed using modified Farnham lysis buffer with syringe passes to assist in mechanical disruption and followed by nuclear lysis in a RIPA variant buffer.

### Sonication conditions offer limited flexibility

Following the optimization of crosslinking, the next critical step is to establish sonication conditions that shear chromatin to an optimal fragment size. Efficient and consistent shearing can lead to higher reproducibility as chromatin length has been shown to affect ChIP-seq quality and sensitivity (Keller et al. 2021). Over-sonication consistently reduces ChIP quality, particularly for transcription factors, while under-sonication can lead to a loss of signal for some targets.

The ChIP-exo 5.0 protocol reported shearing chromatin in IP dilution buffer (containing 0.2% SDS) with a Diagenode Bioruptor for 10 cycles with 30 s on/off intervals to obtain DNA fragments 100 to 500 bp in size. SDS is commonly used to aid sonication, yet it is known to interfere with antibody binding (Qualtiere et al. 1977). It is therefore common for chromatin sonicated in the presence of SDS to be diluted before proceeding to ChIP. To circumvent the use of SDS and to maintain a low volume ChIP reaction, we introduced a chromatin pelleting step and used PBS with protease inhibitors as the sonication buffer. We then systematically evaluated a range of sonication cycles (with 30 s on/off intervals) using a Diagenode Bioruptor Pico and default ‘Easy Shear’ intensity.

As expected, both average fragment size and size distribution decrease with increasing sonication cycles (**Supplementary Figure 4**) and this aligns with decreasing mean insert size between paired-end reads after sequencing (**Figure 4A**). Interestingly, ChIP-exo libraries present acceptable quality metrics after even a single cycle of sonication (**Figure 4A**), even though a smaller proportion of fragments populate the target size range (i.e., 100bp – 500bp) compared with after three sonication cycles (**Supplementary Figure 4**). Key metrics, including FRiP scores and the number of detected CTCF peaks, remained stable for libraries sonicated between one and five cycles but decreased sharply with further sonication, with almost no peaks detected after eight cycles.

**Figure 4.**
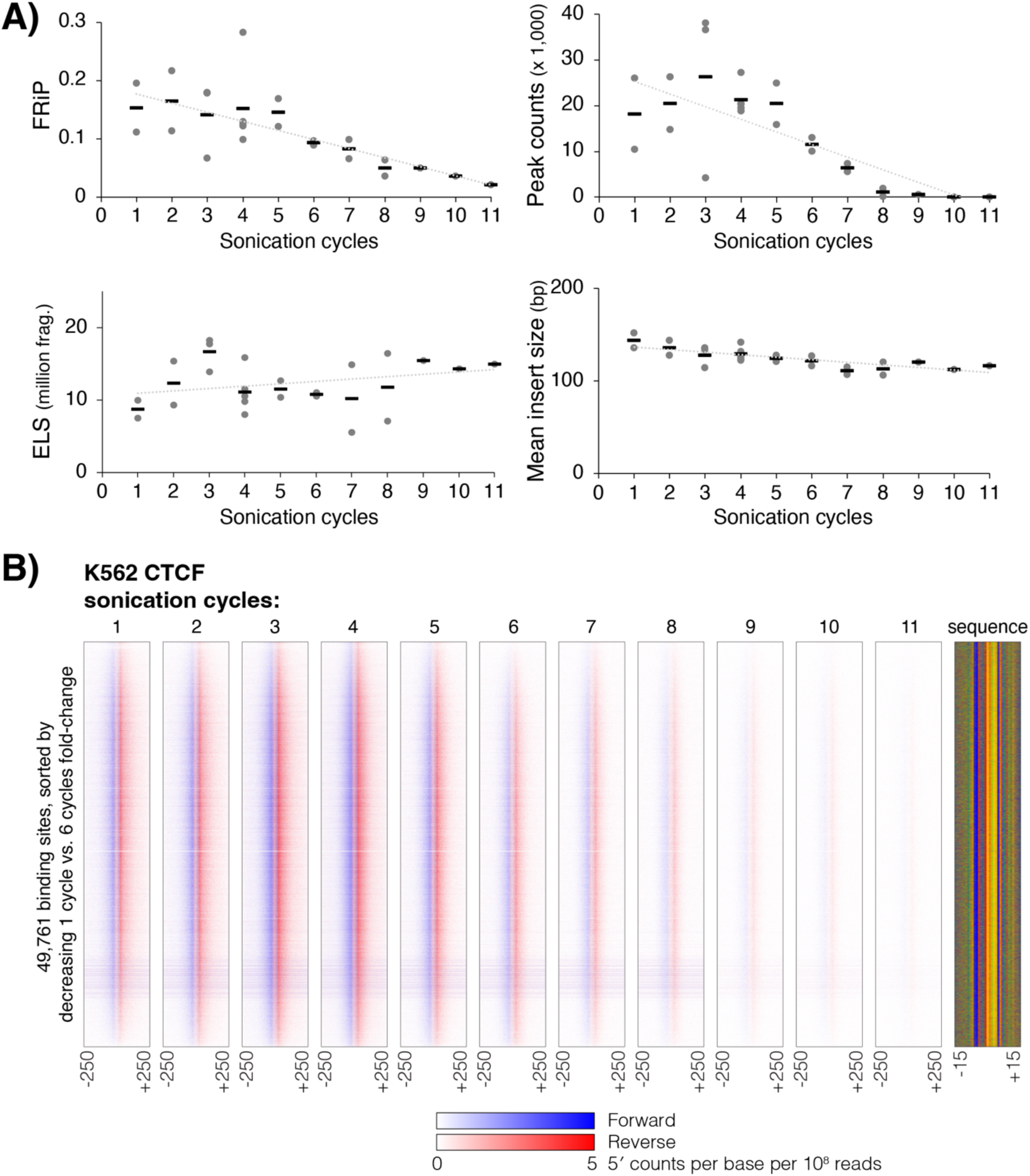
Sonication titrations and the effect of increased shearing on ChIP-exo library quality. **A)** Quality metrics (i.e., FRiP, peak counts, and ELS) and mean insert size (calculated from paired-end sequencing data) are shown for CTCF ChIP-exo libraries prepared with between 1 and 11 cycles of sonication. **B)** Heatmaps display normalized per base enrichment of 5′ read positions from CTCF ChIP-exo libraries prepared with between 1 and 11 cycles of sonication. The heatmaps plot enrichment in 500bp windows centered on all detected CTCF peaks. Blue represents forward strand 5′ enrichment, while red represents reverse strand 5′ enrichment. Binding sites are ordered according to decreasing fold-change enrichment between 1-cycle and 6-cycle libraries.

Interestingly, we found no variation in the identity of CTCF peaks that are detectable with low versus intermediate levels of sonication (**Figure 4B**, **Supplementary Figure 5**). We might expect that higher levels of sonication would “free up” chromatin associated with heterochromatic binding sites, thus leading to a difference in the cohorts of binding sites detected. However, no variation in CTCF enrichment levels beyond that expected between biological replicates was detected when comparing libraries with one and six cycles of sonication (**Supplementary Figure 5**). Rather, a consistent cohort of sites displays ChIP-exo enrichment over a range of sonication cycles, and this signal gradually fades out as sonication cycles are increased (**Figure 4B**).

Thus, while chromatin length affects ChIP-exo quality and sensitivity, there is a range of sonication cycles that yields fragments of useful size and quality, without any impact on the cohorts of binding sites that are detected. Although a range of two to five sonication cycles produced high-quality libraries, we selected four cycles for our protocol due to the reduced variability in peak detection observed across multiple replicates.

Once sheared, chromatin is subjected to immunoprecipitation overnight at 4 °C using an anti-CTCF antibody and Protein A/G magnetic beads, followed by sequential washes to prepare the material for library construction. Our MO-ChIP-exo protocol uses 10 M cells, and 50 μL of Protein A/G Dynabeads following manufacturer recommendations (Invitrogen) with up to 10 μg of antibody.

### Revised library construction reduces sequencing artifacts on patterned flow cells

The initial on-resin steps of library construction, where DNA fragments are prepared for sequencing adapter ligation, required significant modification to address issues arising from the use of an Illumina NextSeq 2000 patterned flow cell sequencer. The ChIP-exo 5.0 protocol involves A-tailing using a Klenow Fragment, followed by Read 2 adapter ligation via TA ligation using T4 DNA ligase and a fill-in reaction using Phi29 polymerase. However, libraries prepared with this method exhibited a high proportion of Read 2 sequences containing poly-G tracts (>10%) and a high rate of undetermined reads containing unexpected barcode combinations (**Figure 5A, 5B**).

**Figure 5.**
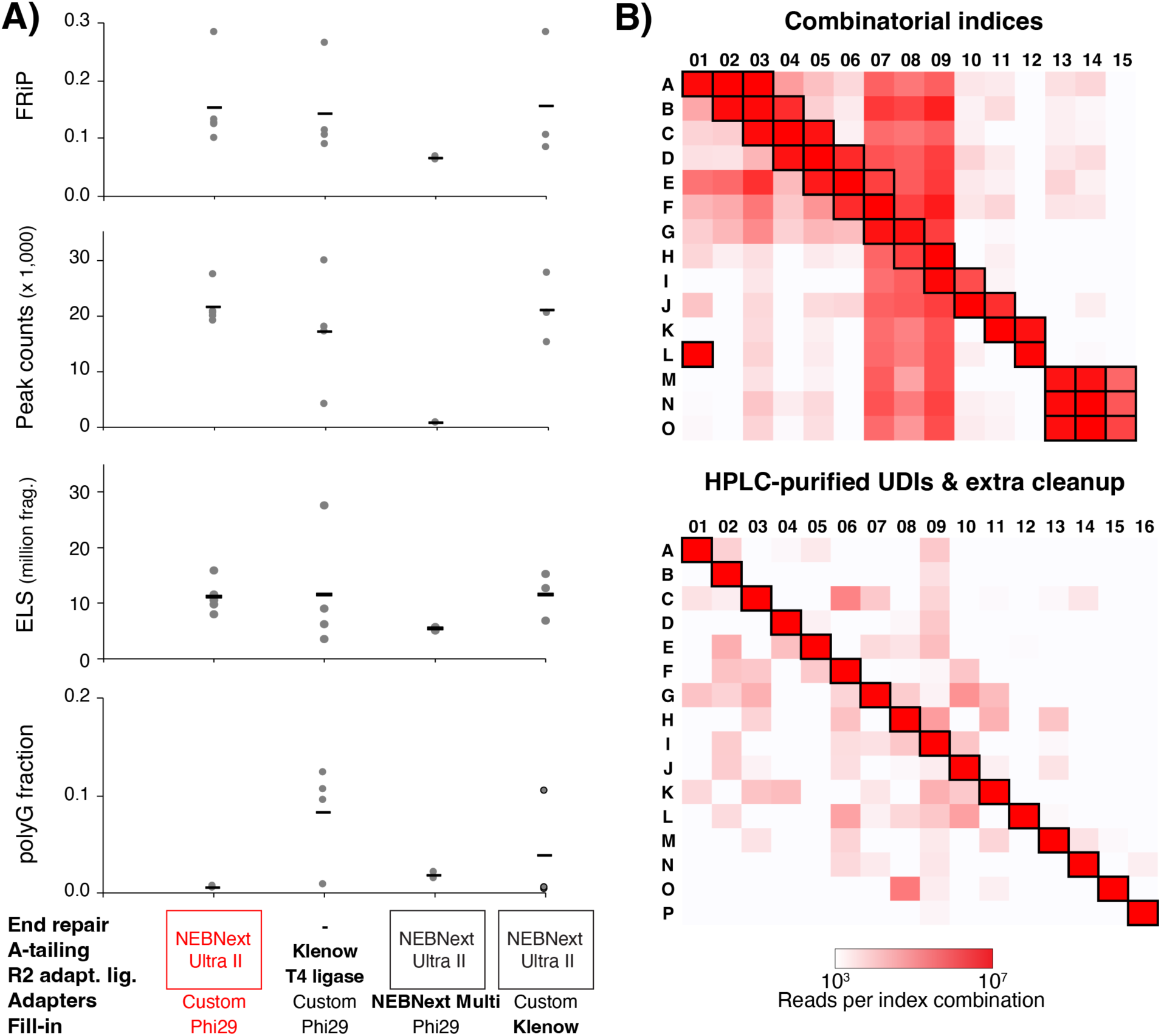
Exploring reaction alternatives confirms the advantage of using a kit to combine end-repair, A-tailing, and first adapter ligation. **A)** Quality metrics are shown for CTCF libraries made using the NEBNext® Ultra™ II DNA Library Prep Kit for the first adapter ligation compared to the custom A-tailing and ligations from ChIP-exo 5.0 (without end-repair), an attempt to use NEBNext Multiplex Oligo adapters, and replacing Phi29 polymerase with Klenow for the fill-in reaction. MO-ChIP-exo conditions are highlighted in red. **B)** Heatmaps summarize the numbers of observed sequencing reads assigned by demultiplexing to each possible combination of read 1 and read 2 indices across two NextSeq 2000 sequencing runs. In each panel, only index combinations whose square is highlighted by a black outline were included in the pooled sequencing library. The top panel summarizes a sequencing run that pooled 38 libraries with combinatorial indices. High read counts are observed outside the expected index combinations, indicating a high rate of index hopping. The bottom panel summarizes a sequencing run that pooled 16 libraries with HPLC-purified unique dual indices and an extra cleanup step after size selection.

The high occurrence of poly-G artifacts, which result from weak or absent signal in the sequencer’s two-color detection chemistry, suggested an inefficient Read 2 adapter ligation reaction. The original ChIP-exo protocol included an end-repair step which was deemed unnecessary in ChIP-exo 5.0 (Rossi et al. 2018). However, the change in sequencing technologies may have exacerbated an existing inefficient ligation issue. To address this, we replaced the custom A-tailing and ligation steps with the NEBNext® Ultra™ II DNA Library Prep Kit for Illumina. This kit combines end-repair and A-tailing into a single polishing reaction followed directly by ligation, streamlining the protocol. Switching to this kit greatly reduced Read 2 poly-G artifacts while maintaining or slightly improving key quality metrics (**Figure 5A**).

We explored additional modifications to the Read 2 adapter ligation step, but these did not improve the results. First, we substituted the custom ChIP-exo adapters with NEBNext® Multiplex Oligos. This change lowered the overall quality metrics (**Figure 5A**) and extended processing times due to the requirement to remove a uracil residue in the adapter’s hairpin loop. Second, we tested an alternative enzyme for the fill-in reaction, which uses Phi29 polymerase to complete the Read 2 adapter sequence. We hypothesized that the Klenow Fragment (3′→ 5′ exo-), which lacks exonuclease activity, might be more effective. However, no difference in library quality was observed between Phi29 and Klenow (**Figure 5A**). Hence, the MO-ChIP-exo protocol retains the use of Phi29 for the fill-in reaction.

The second major issue was the high percentage of undetermined reads, which resulted from using the protocol’s original combinatorial indexing strategy on a patterned flow cell. Specifically, using combinatorial index IDs resulted in a high rate of undetermined reads (averaging ∼19% of sequenced reads per library) when pooled libraries were sequenced on a NextSeq 2000. Patterned flow cell sequencers are known to be more susceptible to index hopping, a phenomenon where reads from one sample are incorrectly assigned to another (Illumina 2018). Analysis of the undetermined reads revealed unexpected index combinations, consistent with index hopping (**Figure 5B, top panel**). To mitigate this issue, we included a second cleanup step using magnetic beads after size selection on the enriched libraries to completely remove any free adapters and possible adapter dimers before pooling libraries. We also switched to using unique dual indices (UDI) that were purified via High-Performance Liquid Chromatography (HPLC). ChIP-exo adapters are >50bp long and incorporating HPLC purification removes truncated sequences that arise during synthesis. The use of UDIs enables bioinformatic filtering of hopped index combinations, which are then correctly classified as undetermined rather than being misassigned. This combination of changes greatly reduced the observed number of hopped indices and the proportion of undetermined reads overall (**Figure 5B**).

Following these critical modifications to the initial Read 2 adapter ligation steps, the remainder of the library construction, including the lambda exonuclease digestion and the Read 1 adapter ligation, proceeds as described in the original ChIP-exo 5.0 protocol (Rossi et al. 2018).

### Refining input quantity, enrichment, and size selection to improve protocol performance

After addressing critical sequencing artifacts, we further refined the protocol to optimize library yield and quality. We first investigated the optimal input quantity. ChIP-exo 5.0 uses 10 M cells per library and reports high quality yields from ∼250,000 cells and detection of CTCF binding signal from as few as 27,000 cells. We tested MO-ChIP-exo with a range of input cell numbers, from 1,000 cells to 50 million (**Figure 6A**). While libraries with comparable complexity can be constructed from 1 M and 5 M cells, their FRiP scores and peaks counts were greatly reduced when compared to the standard 10 M cells (**Figure 6A**). Interestingly, increasing the input to 50 M cells did not improve any of the quality metrics, suggesting that there may be a component of the chromatin immunoprecipitation that is saturated or limiting the input material carried through. Neither increasing the bead volume 4-fold (from 50 μL to 200 μL) nor doubling the concentration of CTCF antibody resulted in improved quality metrics (**Figure 6B**). These results confirmed that our MO-ChIP-exo protocol has achieved a robust balance between cell number, bead volume, and antibody concentration, with 10 M cells representing the optimal input.

**Figure 6.**
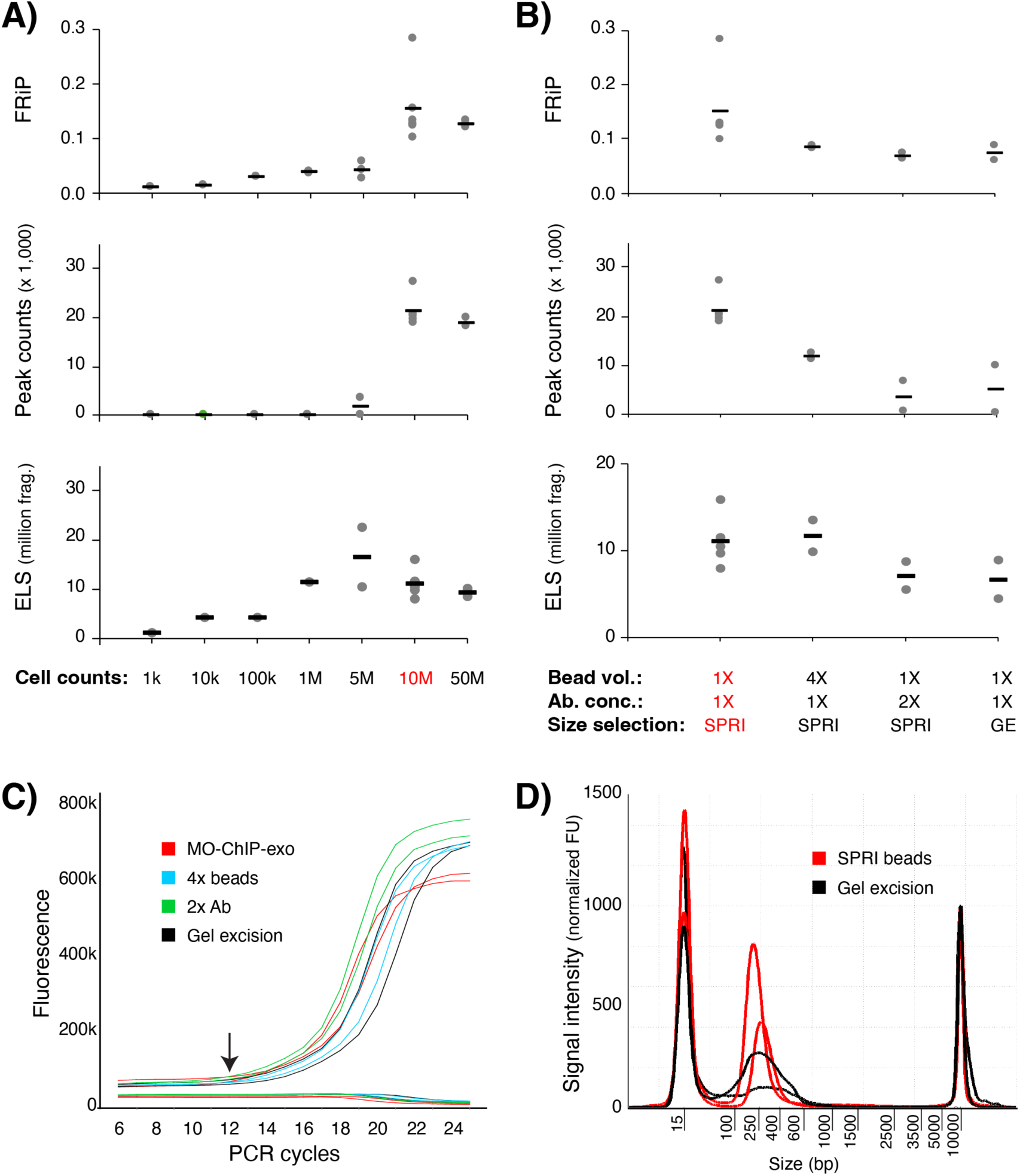
Exploring potential yield/quality improvements: input cell number, antibody and bead proportion, size selection, and enrichment. **A)** Quality metrics for CTCF ChIP-exo libraries when decreasing or increasing the input cell number. **B)** Quality metrics for CTCF ChIP-exo libraries when increasing the Protein A/G beads for antibody capture, doubling the antibody concentration, or using gel excision for library size selection as in ChIP-exo 5.0. **C)** To monitor enrichment, a qPCR was performed using partially amplified libraries (5-cycles) as template, with 7 additional cycles showing sufficient enrichment for completed libraries. **D)** Electropherograms of completed CTCF libraries show a higher yield for libraries size selected with SPRI beads when compared with libraries size-selected using gel excision. MO-ChIP-exo conditions are highlighted throughout in red.

Next, we sought to minimize PCR duplicates by optimizing the library enrichment step. The ChIP-exo 5.0 protocol recommends 18 cycles of PCR amplification, which we observed to produce an increase in sequence duplication rates and an excess of library material (average library concentration of 4 nM with some ranging as high as 18 nM). To avoid over-amplification, we implemented a qPCR-based monitoring strategy, similar to that used for ATAC-seq (Buenrostro et al. 2013). After an initial 5 cycles of PCR, a small aliquot (10%) of the library is monitored for an additional 20 cycles via qPCR using SYBR Green to precisely determine the number of additional cycles necessary to reach the target concentration without reaching saturation (where the target concentration is ∼0.5 nM to allow loading on the NextSeq 2000). This qPCR monitoring strategy reduced total amplification from the original 18 cycles to an average of 12 (i.e., initial 5 plus additional 7), thereby reducing PCR duplicates and increasing the proportion of unique fragments in each library while producing enough fragments for sequencing (average library concentration of 2 nM) (**Figure 6C**).

Finally, we replaced the laborious and lossy gel excision step for size selection with a more efficient magnetic bead-based method. The ChIP-exo 5.0 protocol relied on gel excision (200bp – 500bp) to remove adapter-dimers and to select library fragments of the desired length, but this process is delicate and can lead to sample loss. The ChIP-exo 5.0 protocol states that alternative size selection approaches lead to unacceptably high levels of adapter dimers (Rossi et al. 2018). However, we found that using SPRIselect DNA size selection magnetic beads (Beckman Coulter) for the size selection and cleanup step was faster, required less sample volume, and provided more accurate quantification. Contrary to the original protocol’s concerns, libraries purified with SPRI beads maintained or improved quality metrics, including lower duplication rates, compared to those purified by gel excision (**Figure 6B-6D**). Thus, MO-ChIP-exo incorporates SPRI beads for size selection, in conjunction with TapeStation-based assessment of library fragment distributions. This change streamlined the workflow, reduced processing time, and enhanced overall library quality.

### Validation and application of MO-ChIP-exo

To validate the performance of MO-ChIP-exo, we compared our results to previously published K562 CTCF data from the original (ChIP-exo 1.1) and simplified (ChIP-exo 5.0) protocols, re-analyzing all datasets with our pipeline for consistency. MO-ChIP-exo libraries produce the highest FRiP scores and greatest number of CTCF peaks, with a library complexity comparable to ChIP-exo 1.1 (**Figure 7A**). While ChIP-exo 5.0 libraries exhibited higher ELS scores, this was offset by lower FRiP scores and a significant proportion of undetermined reads (>10%), despite being sequenced in nonpatterned flow cells. Our results suggest that MO-ChIP-exo strikes a balance, preserving beneficial features from the original ChIP-exo 1.1 protocol, such as a polishing step, while integrating new optimizations that improve data quality and streamline the workflow.

**Figure 7.**
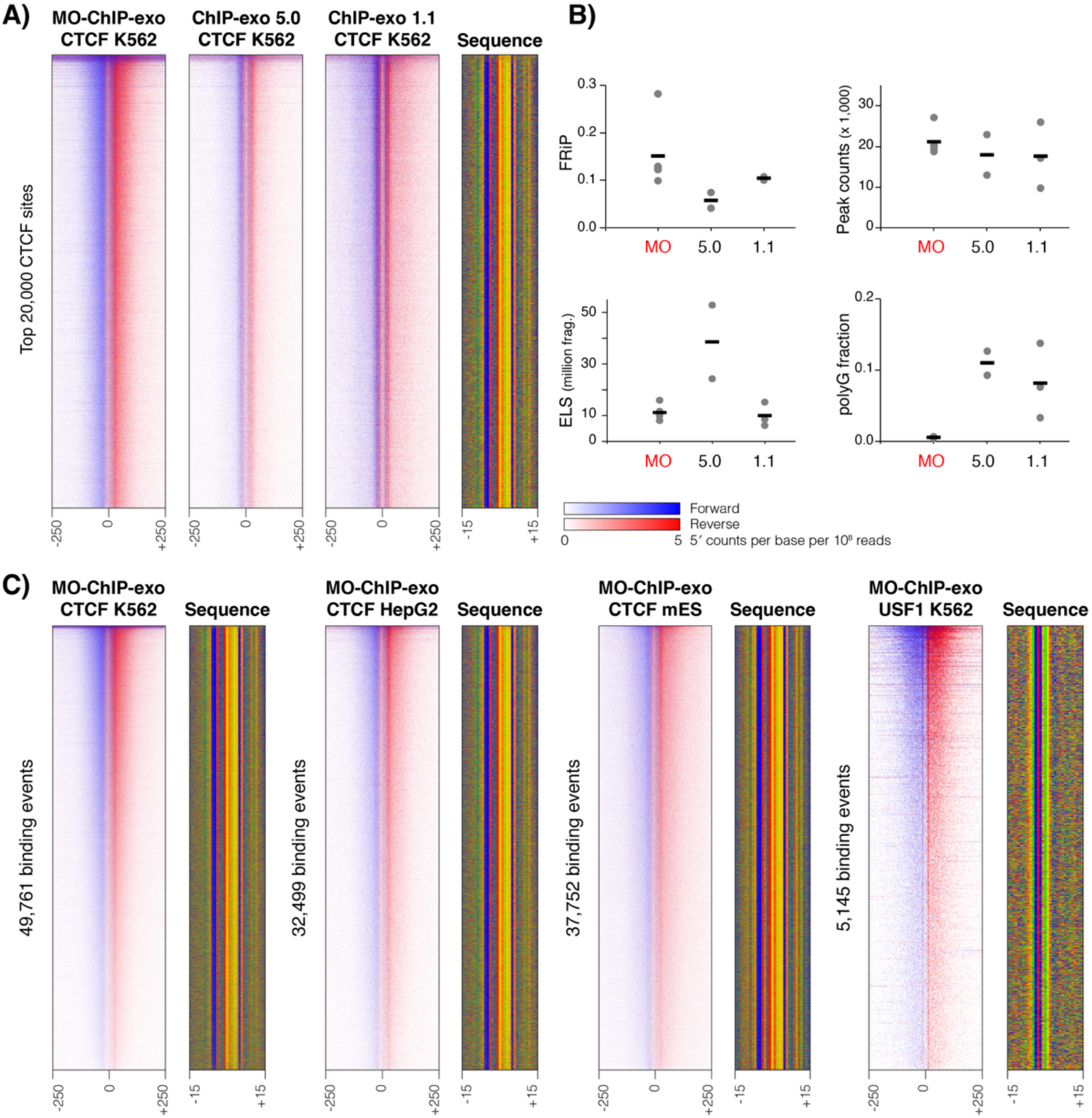
MO-ChIP-exo validation. **A)** MO-ChIP-exo data for CTCF in K562 cells compared with published K562 CTCF data from the simplified ChIP-exo 5.0 and the original ChIP-exo 1.1 protocols (Rossi et al. 2018). Heatmaps display normalized per base enrichment of 5′ read positions from each ChIP-exo protocol. The heatmaps plot enrichment in 500bp windows centered on the top 20,000 CTCF peaks. Blue represents forward strand 5′ enrichment, while red represents reverse strand 5′ enrichment. **B)** Quality metrics (i.e., FRiP, peak counts, and ELS) and percentages of read 2 containing poly-G sequences are shown for MO-ChIP-exo, ChIP-exo 5.0, and ChIP-exo 1.1. **C)** Heatmaps, formatted as in A), displaying MO-ChIP-exo normalized per base enrichment of 5′ read positions across all ChExMix peaks found in K562 CTCF, HepG2 CTCF, mES CTCF, and K562 USF1.

To demonstrate the broader applicability of MO-ChIP-exo, we applied it to characterize CTCF binding in two adherent mammalian cell lines: human HepG2 and mouse embryonic stem cells (mESCs). Both cell lines were trypsinized, quenched, and resuspended in fresh media for crosslinking. Cell counts were obtained at this point to mitigate the effect of cell loss from additional centrifugation steps. While we performed a sonication titration for both cell lines, we determined that the optimal setting of 4 sonication cycles based on K562 cells was also appropriate for these cell lines (**Supplementary Figure 4**). MO-ChIP-exo produced high-quality CTCF ChIP-exo libraries in both cell lines, with quality metrics comparable to the K562 libraries (**Figure 7B**). Finally, we successfully applied the method to target a different transcription factor, USF1, in K562 cells, demonstrating that the MO-ChIP-exo protocol is versatile and not limited to a single protein target (**Figure 7B**).

## DISCUSSION

While ChIP-exo offers unparalleled resolution in mapping protein-DNA interactions, the protocol’s complexity has limited its widespread adoption. This study presents MO-ChIP-exo, a version of the ChIP-exo protocol that is systematically optimized for use in mammalian cells and adapted for compatibility with patterned flow cell sequencing platforms. The main differences between MO-ChIP-exo and the previous ChIP-exo 5.0 protocol are summarized in **Figure 1**. Our work addresses critical, often unreported, methodological variables and introduces modifications that enhance data quality, workflow efficiency, and overall reliability.

One important consideration that has emerged from our results is the crosslinking vehicle. We demonstrate that crosslinking cells directly in their growth media is superior to using PBS. Crosslinking in PBS resulted in lower library quality, fewer detected peaks, and reduced yield of both DNA and the target protein. Our data suggests that this effect is partially due to the loss of cells during the additional centrifugation step to replace media with PBS. The loss of cells can be proportionally minimized if cell harvesting is performed in bulk, but this is not always possible due to cost or sample collection constraints. While using media introduces variability from its components, it avoids sample loss and, in our hands, proved essential for generating high-quality input material. This highlights the importance of empirically determining optimal conditions, as even seemingly minor procedural details can profoundly affect results.

We further optimized the crosslinking reaction itself. Our results show that 1% formaldehyde for 10 minutes produces suitable crosslinking, aligning with other ChIP-seq and ChIP-exo protocols. We elected to use 2-fold molar excess of Tris as a quencher instead of Glycine. Although both are effective, Tris inherently has a greater quenching capacity. We also observed that Glycine significantly lowers the pH of the reaction, which likely contributes to its efficacy by slowing the crosslinking rate. The importance of pH in fixation has been explored for immunofluorescence (Berod et al. 1981), showing that a higher pH achieved optimal fixation. This is in line with our observation that pH may play a part regulating the crosslinking quench. We note that for a robust and abundant target like CTCF, a range of crosslinking conditions may be successful. More stringent crosslinking conditions may be needed for successful detection of less abundant or weaker-binding factors, and we recommend that crosslinking parameters be carefully re-optimized for new targets or epitopes.

Consistent chromatin shearing is also critical, especially for patterned flow cell sequencers like the Illumina NextSeq 2000. These platforms use exclusion amplification, a clustering method that exhibits a bias towards smaller DNA fragments. It is therefore important to balance the generation of smaller fragments, which improves clustering efficiency, with the ability (later in the protocol) to efficiently remove adapter-dimers that can otherwise dominate the sequencing run. ChIP-seq and ChIP-exo protocols typically aim to select DNA fragments between 100 and 500bp in length. Our sonication titrations show that a high proportion of DNA fragments will fall in this optimal size range after one to five sonication cycles. We selected four cycles as the standard for MO-ChIP-exo, as it consistently produced the highest number of peaks with low replicate variability. This number of cycles also provided consistent results across several cell lines.

The most significant modifications in our protocol were driven by the need to resolve sequencing artifacts specific to patterned flow cells. Our initial libraries were plagued by a high rate of index hopping and the presence of poly-G tracts in Read 2. We addressed index hopping by introducing a second magnetic bead-based cleanup step after size selection to remove free adapters. We also replaced the original combinatorial indices with HPLC-purified UDIs, which allow for the bioinformatic removal of misassigned reads. The poly-G artifact, which pointed to an inefficient first adapter ligation, was resolved by re-introducing an end-repair step. By incorporating a modern library preparation kit, we consolidated the end-repair, A-tailing, and ligation reactions, which not only fixed the ligation issue but also streamlined the protocol by eliminating intermediate washes.

Finally, we refined several steps to improve workflow efficiency and yield. We confirmed that samples with 10 million cells provide the optimal input, as lower cell numbers reduced data quality and higher cell numbers offered no benefit. To minimize PCR duplicates, we replaced the fixed-cycle enrichment with a qPCR-based monitoring approach that significantly reduces total library amplification. Furthermore, we substituted the gel excision step for size selection with a faster and more efficient magnetic bead-based method, which improved DNA recovery and lowered duplication rates.

ChIP-exo is a laborious protocol with numerous steps that are critical to overall success. A major challenge is the lack of informative quality control checkpoints that assess the efficiency of enzymatic reactions, making troubleshooting very difficult before sequencing. ChIP-exo’s signature exonuclease step degrades about half of an already limited amount of starting material. While we recommend integrating the use of a fragment analyzer to assess library quality and fragment size distribution, these DNA library metrics are not predictive of the success of the ChIP reaction. In our investigations, for example, estimated library size was not a good predictor of FRiP score or peak counts (**Supplementary Figure 6**). The MO-ChIP-exo protocol was designed to navigate these difficulties, providing a more robust and reliable path to generating high-quality libraries. Combined with the critical updates for sequencer compatibility, our improvements result in a validated protocol that produces libraries with superior FRiP scores and peak counts compared to previous ChIP-exo protocols. MO-ChIP-exo has been successfully applied to different cell types and alternative transcription factors, demonstrating its versatility.

## MATERIALS AND METHODS

### Cell culture, Crosslinking and Harvest

K562 (CCL-243) and HepG2 (HB-8065) cells were obtained from the American Type Culture Collection (ATCC) handled and sub-cultured following the manufacturers recommendations. A17 mES cells were cultured as previously described (Bulajić et al. 2020). K562 suspension cells were counted and directly crosslinked in media or pelleted and resuspended for crosslinking in PBS. Adherent cells were washed with PBS, trypsinized, quenched and resuspended in media or PBS for counting and crosslinking. All cell counts were performed with a trypan blue dye exclusion test using a hemocytometer. For MO-ChIP-exo, 10 min of crosslinking with 1% formaldehyde was quenched with a 3M Tris pH 7.8. Variations of the crosslinking time, formaldehyde concentration and quencher were used as reported. Crosslinked cells were washed with PBS + 1X CPI and flash frozen in liquid nitrogen. A detailed crosslinking and harvest protocol is provided in **Supplementary Methods**.

### MO-ChIP-exo Sonication and Library Construction

Crosslinked cell pellets (10 M cells per reaction) were subjected to cell and nuclear lysis. Lysates were resuspended in PBS + 1X CPI for sonication in a Diagenode Bioruptor Pico. The ‘Go & Shear’ setting was used with the validated parameters on the ‘Easy Mode’ for 30’’ON/OF with increasing cycles (1-11 as described). An aliquot of 15 μL (∼0.5 M cells) was used to reverse crosslink and purify the DNA for a sonication assessment using the Agilent Tape Station D5000 kit. The sonicated crosslinked chromatin was pre-cleared before ChIP using high speed centrifugation (14k rpm at 4°C for 15 min). ChIP antibodies [Anti-CTCF (Sigma 07-729), Anti-USF1-1B8 (DSHB USF1-1B8-6/13/19), IgG (Sigma i5006), Anti-GATA1 (Santa Cruz Biotechnology SC-266), Anti-FoxA1 (Santa Cruz Biotechnology SC-101058), Anti-SOX2 (R&D AF2018)] were pre-conjugated to 50 μL Protein A or G Dynabeads (Invitrogen) before an overnight incubation with the antibody of interest at 4°C. Bead-bound chromatin was washed and subjected to simultaneous End-repair and A-tailing with a subsequent TA ligation of the first HPLC-purified adapter using the NEBNext® Ultra™ II DNA Library Prep Kit. After washes, a fill-in reaction was performed using Phi29 DNA Polymerase (NEB). This reaction was followed by another wash and Lambda Exonuclease (NEB) digestion. Reverse crosslinking was performed overnight to release the DNA. Released DNA was purified using Mag-Bind® TotalPure NGS beads (Omega Bio-Tek) before ligating the 2^nd^ HPLC-purified adapter using T4 DNA ligase (Qiagen). A DNA cleanup was performed using Mag-Bind® TotalPure NGS beads before PCR enrichment for 12 cycles. This was followed by size selection using SPRI select beads (Beckman Coulter) and an additional DNA cleanup to remove excess adapters. Completed libraries were analyzed in the Tape Station using High Sensitivity D5000 ScreenTape and Reagents. Finally, libraries were individually quantified through qPCR using NEBNext® Library Quantification Kit, For Illumina® (NEB) in an ABI StepOne Plus Real Time PCR System. A detailed MO-ChIP-exo protocol is provided in **Supplementary Methods**.

### Input assessment using ChIP-qPCR and Western blot

The input and CTCF-ChIPed DNA from different crosslinking conditions were quantified using a Qubit™ 1X dsDNA HS Assay Kit (Thermo Scientific) and used as templates for qPCR comparative enrichment using the % input method. Anti-CTCF (Sigma 07-729) was used for the ChIP. The NEB Luna® Universal qPCR Master Mix was used with primers targeting the KLK locus (KLK2, KLK5) and a CTCF unbound region (WNT2) (Khoury et al. 2020). CTCF-ChIPed DNA was normalized to input and each condition was quantified in triplicate. To assess CTCF protein abundance, Western blot was used. To prepare lysates, K562 cells were washed in ice-cold PBS + 1XCPI and collected by centrifugation. The cell pellet was lysed in 1.5X Laemmli buffer and boiled at 95 °C for 5-10 minutes. After centrifugation at 14k rpm for 10min, the supernatant was collected and stored at -20°C. Lysates were re-boiled for 3min upon thawing and separated through SDS-PAGE using 8% polyacrylamide gels. Gels were transferred using the Trans-Blot Turbo Transfer System and RTA Mini 0.2 μm Nitrocellulose Transfer Kit (Biorad). Blots were blocked in 5% milk in TBS-T for 1 hr at RT. CTCF (G-8) HRP (Santa Cruz Biotechnology sc-271474) was diluted 1:1,000 in 5% milk in TBS-T and incubated overnight with gentle rocking at 4°C. TBS-T washes were performed before developing with Pierce™ ECL Western Blotting Substrate (Life Technologies) using a ChemiDoc Imaging System (Biorad). After imaging, the blot was stripped and re-probed using Beta Tubulin (BT7R) (Life Technologies MA5-16308-HRP) 1:10,000 as a loading control.

### Monitoring qPCR

MO-ChIP-exo library PCR amplification using Phusion High Fidelity DNA Polymerase (Thermo Scientific) was paused after 5-cycles in a subset of libraries. A 5 μL aliquot was removed from each library and used as template for qPCR. Library aliquots were amplified for another 20-cycles in an ABI Step One System adding SYBR green and ROX to the master mix as previously described (Buenrostro et al. 2013). A multicomponent plot (fluorescent signal vs cycle number) was obtained and used to determine the number of additional cycles necessary to reach optimal library concentration for sequencing. The number of cycles before the curve starts to increase exponentially was chosen as optimal. Libraries paused at 5-cycles were enriched for an additional 7-cycles, size selected, cleaned up, and analyzed in the Tape Station using High Sensitivity D5000 ScreenTape and Reagents. Completed libraries were quantified using the Library Quantification Kit, For Illumina® (NEB) to confirm sufficient library for sequencing.

### pH monitoring

Changes in pH were monitored during crosslinking in multiple crosslinking and quenching conditions using an Orion™ Star™ A215 pH/Conductivity Benchtop Multiparameter Meter (Thermo Scientific). An aliquot ∼25 mL of media (IMDM + 10% FBS) or PBS was used to measure pH before and after adding formaldehyde to 1% final concentration. After 5-10min a quencher was added, and the pH was re-measured.

### Sequencing

Completed libraries were pooled for each sequencing run based on their concentration and read allocation with a 1% PhiX spike-in, treated with Illumina’s Free Adapter Blocking Reagent and cleaned up using Mag-Bind® TotalPure NGS beads. Final pool concentration was determined through qPCR using NEB Next® Library Quantification Kit, For Illumina® in an ABI StepOne Plus Real Time PCR System before diluting to 0.65 nM for loading. An Illumina NextSeq 2000 Sequencing System was used with a NextSeq™ 1000/2000 P2 XLEAP-SBS™ Reagent Kit (100 Cycle) for paired-end sequencing with 50 bp each for read1 and read2.

### Data Analysis

#### Read alignment

Adapter sequences were first trimmed from the paired-end fastq files using fastp (v.0.23.2) (S. Chen et al. 2018). Paired-end reads were aligned using BWA-MEM (v.0.7.17) (Li and Durbin 2010) with parameter -M against the human (hg38) or mouse (mm10) genomes. BAM files were sorted using samtools sort (v.1.16.1) (Li et al. 2009) and duplicate read pairs were marked using Picard MarkDuplicates (v.2.24.1) with arguments REMOVE_DUPLICATES=’false’ ASSUME_SORTED=’true’ DUPLICATE_SCORING_STRATEGY=’SUM_OF_BASE_QUALITIES’ READ_NAME_REGEX=’[a-zA-Z0-9]+:[0-9]:([0-9]+):([0-9]+):([0-9]+).*.’. BAM files were then filtered to remove reads that are duplicated, unmapped, or not in proper pairs using samtools view (v.1.16.1) with arguments -h -b -f 0×3 -F 0×404.

#### Peak-finding

Peaks for each individual ChIP-exo sample were determined using ChExMix (v0.52) (Yamada et al. 2018) using the following parameters: --threads 4 --expt ${expt} --ctrl ${ctrl} -- format BAM --scalewin 1000 --noread2 --round 3 --minmodelupdateevents 50 --prlogconf -4 -- alphascale 1.0 --betascale 0.05 --epsilonscale 0.2 --minroc 0.7 --minmodelupdaterefs 25 --pref - --numcomps 500 --win 250 --back ${back} --memepath ${memePath} --mememinw 6 --mememaxw 18 --seq ${genome} --exclude ${exclude} --q 0.01 --minfold 1.5.

Blacklist files (the ${exclude} argument) were sourced from ENCODE. Background files (the ${back} argument) are 2^nd^-order Markov models created from the human or mouse genomes. Control samples (the ${ctrl} argument) represent pooled BAM files of all relevant control samples in each cell type, merged using samtools merge (v.1.16.1) (Li et al. 2009). This merger was performed across 13 K562 control samples (totaling 40,362,133 read pairs), 5 HepG2 control samples (13,990,398 read pairs), and 14 mES control samples (93,023,738 read pairs).

#### FRiP scores

Fraction of Reads in Peaks (FRiP) scores for each sample are calculated by counting the fraction of mapped read1 reads (i.e., the ChIP-exo resolution read) that intersect with peaks according to bedtools intersect (v2.30.0) (Quinlan and Hall 2010). FRiP scores are typically calculated using peaks called from a given sample. However, the number of peaks called in each dataset depends not only on the quality of the ChIP experiment, but also on other parameters such as sequencing depth and library complexity. To ensure that FRiP scores are a comparable metric of quality across samples, we use a common set of peaks for each TF and cell type. The FRiP scores presented in the manuscript were defined with respect to peaks determined from ENCODE ChIP-seq data for each relevant TF and cell type. Specifically, we combined the following ENCODE narrowPeaks BED files using bedtools sort and bedtools merge (v2.30.0):

- K562 CTCF: ENCFF221SKA, ENCFF582SNT, ENCFF660GHM, ENCFF736NYC, ENCFF769AUF
- K562 USF1: ENCFF255QDL
- HepG2 CTCF: ENCFF119PKI, ENCFF199YFA, ENCFF205OKL, ENCFF240MUS, ENCFF664UGR
- mES CTCF: ENCFF052KVO, ENCFF533APC

We also tested an alternative strategy where peaks for each TF and cell type were determined by performing peak-finding on merged sets of samples that each individually yield more than 1,000 peaks. This approach yielded FRiP scores that are highly correlated with FRiP scores from ENCODE-defined peaks (Pearson correlation coefficient >0.99 for each TF/cell type combination) and essentially unchanged relative rankings (Spearman rank correlation >0.99).

#### Estimated Library Size (ELS)

We calculate the estimated complexity of a ChIP-exo DNA library (i.e., the number of unique DNA fragments in the initial library) using the same procedure as implemented in the EstimateLibraryComplexity function of the Picard toolkit (*Picard Toolkit* 2019). Specifically, this procedure is based on the following equation:

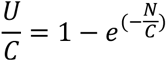

where *C* is the number of distinct molecules in the library, *N* is the number of paired-end reads (total mapped paired-end reads in our usage), and *U* is the number of unique paired-end reads observed (number of deduplicated mapped paired-end reads in our usage). The value of *C* is solved using a bisection search. Note that this framing of the problem assumes that the sequencing process is an unbiased sampling with replacement, governed by a Poisson distribution. In reality, PCR and sequencing introduce substantial biases, so our library size estimates are unlikely to be highly accurate. However, the calculated ELS values should be appropriate for comparing the relative library complexity across experimental conditions, as they are used here.

#### Poly-G read rates

Poly-G-containing reads are defined as those containing 10 or more consecutive Gs. The rates were calculated using an awk script applied to the fastq files.

#### Insert sizes

Read insert size distributions were calculated using Picard CollectInsertSizeMetrics (v.2.24.1) (*Picard Toolkit* 2019).

#### Differential binding analysis

To assess whether CTCF binding varied according to sonication levels, we compared CTCF ChIP-exo enrichment between samples prepared with 1 sonication cycle and 6 sonication cycles (**Figure 4B**, **Supplementary Figure 4**). The 6-cycle experiment was chosen because ChIP-exo signal is too weak in higher cycle experiments. ChIP-exo reads overlapping 200bp windows centered on CTCF binding sites were counted for each replicate in each condition. EdgeR (v.4.4.2) (Robinson et al. 2010) was then used to normalize and compare enrichment levels across conditions.

## Supporting information

Supplemental Protocol

## DATA AVAILABILITY

All raw and processed sequencing data are available from GEO accession number GSE305217.

## ACKNOWLEDGEMENTS

This study was supported by the National Institutes of Health grant R35GM144135. We thank Dr. Cheryl Keller, Dr. William Lai, Dr. Joe Reese, and members of the Center for Eukaryotic Gene Regulation for valuable discussions during our optimization of the ChIP-exo protocol.

## REFERENCES

Abbasova, Leyla, Paulina Urbanaviciute, Di Hu, et al. 2025. “CUT&Tag Recovers up to Half of ENCODE ChIP-Seq Histone Acetylation Peaks.” Nature Communications 16 (1): 2993. 10.1038/s41467-025-58137-2.

Akerström, B, and L Björck. 1986. “A Physicochemical Study of Protein G, a Molecule with Unique Immunoglobulin G-Binding Properties.” Journal of Biological Chemistry 261 (22): 10240–47. 10.1016/S0021-9258(18)67515-5.

Albert, Istvan, Travis N. Mavrich, Lynn P. Tomsho, et al. 2007. “Translational and Rotational Settings of H2A.Z Nucleosomes across the Saccharomyces Cerevisiae Genome.” Nature 446 (7135): 572–76. 10.1038/nature05632.

Baranello, Laura, Fedor Kouzine, Suzanne Sanford, and David Levens. 2016. “ChIP Bias as a Function of Cross-Linking Time.” Chromosome Research 24 (2): 175–81. 10.1007/s10577-015-9509-1.

Berod, A., B. K. Hartman, and J. F. Pujol. 1981. “Importance of Fixation in Immunohistochemistry: Use of Formaldehyde Solutions at Variable pH for the Localization of Tyrosine Hydroxylase.” The Journal of Histochemistry and Cytochemistry: Official Journal of the Histochemistry Society 29 (7): 844–50. 10.1177/29.7.6167611.

Buenrostro, Jason D., Paul G. Giresi, Lisa C. Zaba, Howard Y. Chang, and William J. Greenleaf. 2013. “Transposition of Native Chromatin for Fast and Sensitive Epigenomic Profiling of Open Chromatin, DNA-Binding Proteins and Nucleosome Position.” Nature Methods 10 (12): 1213–18. 10.1038/nmeth.2688.

Bulajić, Milica, Divyanshi Srivastava, Jeremy S. Dasen, Hynek Wichterle, Shaun Mahony, and Esteban O. Mazzoni. 2020. “Differential Abilities to Engage Inaccessible Chromatin Diversify Vertebrate Hox Binding Patterns.” *Development (Cambridge*, England) 147 (22): dev194761. 10.1242/dev.194761.

Chen, Hebing, Yao Tian, Wenjie Shu, Xiaochen Bo, and Shengqi Wang. 2012. “Comprehensive Identification and Annotation of Cell Type-Specific and Ubiquitous CTCF-Binding Sites in the Human Genome.” PLoS ONE 7 (7): e41374. 10.1371/journal.pone.0041374.

Chen, Shifu, Yanqing Zhou, Yaru Chen, and Jia Gu. 2018. “Fastp: An Ultra-Fast All-in-One FASTQ Preprocessor.” Bioinformatics 34 (17): i884–90. 10.1093/bioinformatics/bty560.

Euskirchen, Ghia M., Joel S. Rozowsky, Chia-Lin Wei, et al. 2007. “Mapping of Transcription Factor Binding Regions in Mammalian Cells by ChIP: Comparison of Array- and Sequencing-Based Technologies.” Genome Research 17 (6): 898–909. 10.1101/gr.5583007.

Feingold, E. A., P. J. Good, M. S. Guyer, S. Kamholz, and, et al. 2004. “The ENCODE (ENCyclopedia Of DNA Elements) Project.” SPECIAL SECTION: GENES IN ACTION. *Science* (Washington, United States) 306 (5696): 636–40.

Gilmour, D S, and J T Lis. 1984. “Detecting Protein-DNA Interactions in Vivo: Distribution of RNA Polymerase on Specific Bacterial Genes.” Proceedings of the National Academy of Sciences of the United States of America 81 (14): 4275–79. 10.1073/pnas.81.14.4275.

Giresi, Paul G., Jonghwan Kim, Ryan M. McDaniell, Vishwanath R. Iyer, and Jason D. Lieb. 2007. “FAIRE (Formaldehyde-Assisted Isolation of Regulatory Elements) Isolates Active Regulatory Elements from Human Chromatin.” Genome Research 17 (6): 877–85. 10.1101/gr.5533506.

He, Qiye, Jeff Johnston, and Julia Zeitlinger. 2015. “ChIP-Nexus Enables Improved Detection of in Vivo Transcription Factor Binding Footprints.” Nature Biotechnology 33 (4): 395–401. 10.1038/nbt.3121.

Hoffman, Elizabeth A., Brian L. Frey, Lloyd M. Smith, and David T. Auble. 2015. “Formaldehyde Crosslinking: A Tool for the Study of Chromatin Complexes.” The Journal of Biological Chemistry 290 (44): 26404–11. 10.1074/jbc.R115.651679.

Illumina. 2018. “Effects of Index Misassignment on Multiplexing and Downstream Analysis.” 2018 770-2017-004-D | 1. https://www.illumina.com/content/dam/illumina-marketing/documents/products/whitepapers/index-hopping-white-paper-770-2017-004.pdf?linkId=36607862.

Ji, Hongjie, Zuhua Qiu, Yuzhuo Wang, et al. 2022. “The Effect of Crosslinking Concentration, Time, Temperature and pH on the Characteristic of Genipin-Crosslinked Small Intestinal Submucosa.” Materials Today Communications 33 (December): 104482. 10.1016/j.mtcomm.2022.104482.

Johnson, David S., Ali Mortazavi, Richard M. Myers, and Barbara Wold. 2007. “Genome-Wide Mapping of in Vivo Protein-DNA Interactions.” Science 316 (5830): 1497–502. 10.1126/science.1141319.

Kasinathan, Sivakanthan, Guillermo A. Orsi, Gabriel E. Zentner, Kami Ahmad, and Steven Henikoff. 2014. “High-Resolution Mapping of Transcription Factor Binding Sites on Native Chromatin.” Nature Methods 11 (2): 203–9. 10.1038/nmeth.2766.

Kaya-Okur, Hatice S., Steven J. Wu, Christine A. Codomo, et al. 2019. “CUT&Tag for Efficient Epigenomic Profiling of Small Samples and Single Cells.” Nature Communications 10(1): 1930. 10.1038/s41467-019-09982-5.

Keller, Cheryl A., Alexander Q. Wixom, Elisabeth F. Heuston, et al. 2021. “Effects of Sheared Chromatin Length on ChIP-Seq Quality and Sensitivity.” G3 (Bethesda, Md.) 11 (6): jkab101. 10.1093/g3journal/jkab101.

Khoury, Amanda, Joanna Achinger-Kawecka, Saul A. Bert, et al. 2020. “Constitutively Bound CTCF Sites Maintain 3D Chromatin Architecture and Long-Range Epigenetically Regulated Domains.” Nature Communications 11 (1): 54. 10.1038/s41467-019-13753-7.

Kim, Somi, Nam-Kyung Yu, and Bong-Kiun Kaang. 2015. “CTCF as a Multifunctional Protein in Genome Regulation and Gene Expression.” Experimental & Molecular Medicine 47 (6): e166–e166. 10.1038/emm.2015.33.

Lai, William K. M., Luca Mariani, Gerson Rothschild, et al. 2021. “A ChIP-Exo Screen of 887 PCRP Transcription Factor Antibodies in Human Cells.” Preprint, bioRxiv, March 3. 10.1101/2020.06.08.140046.

Landt, Stephen G., Georgi K. Marinov, Anshul Kundaje, et al. 2012. “ChIP-Seq Guidelines and Practices of the ENCODE and modENCODE Consortia.” Genome Research 22 (9): 1813–31. 10.1101/gr.136184.111.

Li, Heng, and Richard Durbin. 2010. “Fast and Accurate Long Read Alignment with Burrows-Wheeler Transform.” Bioinformatics 26 (5): 589–95. 10.1093/bioinformatics/btp698.

Li, Heng, Bob Handsaker, Alec Wysoker, et al. 2009. “The Sequence Alignment/Map Format and SAMtools.” Bioinformatics 25 (16): 2078–79. 10.1093/bioinformatics/btp352.

Moore, Jill E., Michael J. Purcaro, Henry E. Pratt, et al. 2020. “Expanded Encyclopaedias of DNA Elements in the Human and Mouse Genomes.” Nature 583 (7818): 699–710. 10.1038/s41586-020-2493-4.

Park, Brandon J., Shan Hua, Karli D. Casler, et al. 2025. “CUT&Tag Identifies Repetitive Genomic Loci That Are Excluded from ChIP Assays.” Preprint, bioRxiv, February 5. 10.1101/2025.02.03.636299.

Picard Toolkit. 2019. GitHub Repository. Broad Institute. https://broadinstitute.github.io/picard/.

Qualtiere, L. F., A. G. Anderson, and P. Meyers. 1977. “Effects of Ionic and Nonionic Detergents on Antigen-Antibody Reactions.” Journal of Immunology (Baltimore, Md.: 1950) 119 (5): 1645–51.

Quinlan, Aaron R., and Ira M. Hall. 2010. “BEDTools: A Flexible Suite of Utilities for Comparing Genomic Features.” *Bioinformatics (Oxford*, England) 26 (6): 841–42. 10.1093/bioinformatics/btq033.

Quon, Sara, Bingfei Yu, Brendan E. Russ, et al. 2023. “DNA Architectural Protein CTCF Facilitates Subset-Specific Chromatin Interactions to Limit the Formation of Memory CD8+ T Cells.” Immunity 56 (5): 959–978.e10. 10.1016/j.immuni.2023.03.017.

Rhee, Ho Sung, Alain R. Bataille, Liye Zhang, and B. Franklin Pugh. 2014. “Subnucleosomal Structures and Nucleosome Asymmetry across a Genome.” Cell 159 (6): 1377–88. 10.1016/j.cell.2014.10.054.

Rhee, Ho Sung, and B. Franklin Pugh. 2011. “Comprehensive Genome-Wide Protein-DNA Interactions Detected at Single Nucleotide Resolution.” Cell 147 (6): 1408–19. 10.1016/j.cell.2011.11.013.

Rhee, Ho Sung, and B. Franklin Pugh. 2012. “ChIP-exo Method for Identifying Genomic Location of DNA-Binding Proteins with Near-Single-Nucleotide Accuracy.” Current Protocols in Molecular Biology 100 (1). 10.1002/0471142727.mb2124s100.

Robinson, Mark D., Davis J. McCarthy, and Gordon K. Smyth. 2010. “edgeR: A Bioconductor Package for Differential Expression Analysis of Digital Gene Expression Data.” Bioinformatics 26 (1): 139–40. 10.1093/bioinformatics/btp616.

Rossi, Matthew J., Prashant K. Kuntala, William K. M. Lai, et al. 2021. “A High-Resolution Protein Architecture of the Budding Yeast Genome.” Nature 592 (7853): 309–14. 10.1038/s41586-021-03314-8.

Rossi, Matthew J., William K. M. Lai, and B. Franklin Pugh. 2018. “Simplified ChIP-Exo Assays.” Nature Communications 9 (1): 2842. 10.1038/s41467-018-05265-7.

Rozbeský, Daniel, Michal Rosůlek, Zdeněk Kukačka, Josef Chmelík, Petr Man, and Petr Novák. 2018. “Impact of Chemical Cross-Linking on Protein Structure and Function.” Analytical Chemistry 90 (2): 1104–13. 10.1021/acs.analchem.7b02863.

Schroeder, Barbara, Hoa Le Xuan, Jule L. Völzke, and Michael G. Weller. 2019. “Preactivation Crosslinking—An Efficient Method for the Oriented Immobilization of Antibodies.” Methods and Protocols 2 (2): 35. 10.3390/mps2020035.

Serandour, Aurelien A., Gordon D. Brown, Joshua D. Cohen, and Jason S. Carroll. 2013. “Development of an Illumina-Based ChIP-Exonuclease Method Provides Insight into FoxA1-DNA Binding Properties.” Genome Biology 14 (12): R147. 10.1186/gb-2013-14-12-r147.

Skene, Peter J., and Steven Henikoff. 2018. CUT&RUN: Targeted in Situ Genome-Wide Profiling with High Efficiency for Low Cell Numbers. January 16. https://www.protocols.io/view/cut-run-targeted-in-situ-genome-wide-profiling-wit-mgjc3un.

Sutherland, Brent W., Judy Toews, and Juergen Kast. 2008. “Utility of Formaldehyde Cross-Linking and Mass Spectrometry in the Study of Protein–Protein Interactions.” Journal of Mass Spectrometry 43 (6): 699–715. 10.1002/jms.1415.

Watanabe, Hideki, Hiroyuki Matsumaru, Ayako Ooishi, et al. 2009. “Optimizing pH Response of Affinity between Protein G and IgG Fc: HOW ELECTROSTATIC MODULATIONS AFFECT PROTEIN-PROTEIN INTERACTIONS*.” Journal of Biological Chemistry 284 (18): 12373–83. 10.1074/jbc.M809236200.

Xu, Bingxiang, Xiaomeng Gao, Xiaoli Li, Feifei Li, and Zhihua Zhang. 2024. “Crosslinking Intensity Modulates the Reliability and Sensitivity of Chromatin Conformation Detection at Different Structural Levels.” Communications Biology 7 (1): 1–13. 10.1038/s42003-024-06904-0.

Yamada, Naomi, Prashant Kumar Kuntala, B. Franklin Pugh, and Shaun Mahony. 2020. “ChExMix: A Method for Identifying and Classifying Protein-DNA Interaction Subtypes.” Journal of Computational Biology: A Journal of Computational Molecular Cell Biology 27(3): 429–35. 10.1089/cmb.2019.0466.

Yamada, Naomi, William K. M. Lai, Nina Farrell, B. Franklin Pugh, and Shaun Mahony. 2018. “Characterizing Protein-DNA Binding Event Subtypes in ChIP-Exo Data.” Bioinformatics 35 (6): 903–13. 10.1093/bioinformatics/bty703.

Yen, Kuangyu, Vinesh Vinayachandran, and B. Franklin Pugh. 2013. “SWR-C and INO80 Chromatin Remodelers Recognize Nucleosome-Free Regions near +1 Nucleosomes.” Cell 154 (6): 1246–56. 10.1016/j.cell.2013.08.043.

