## Supplemental Figures for "Optimized ChIP-exo for mammalian cells and patterned sequencing flow cells"

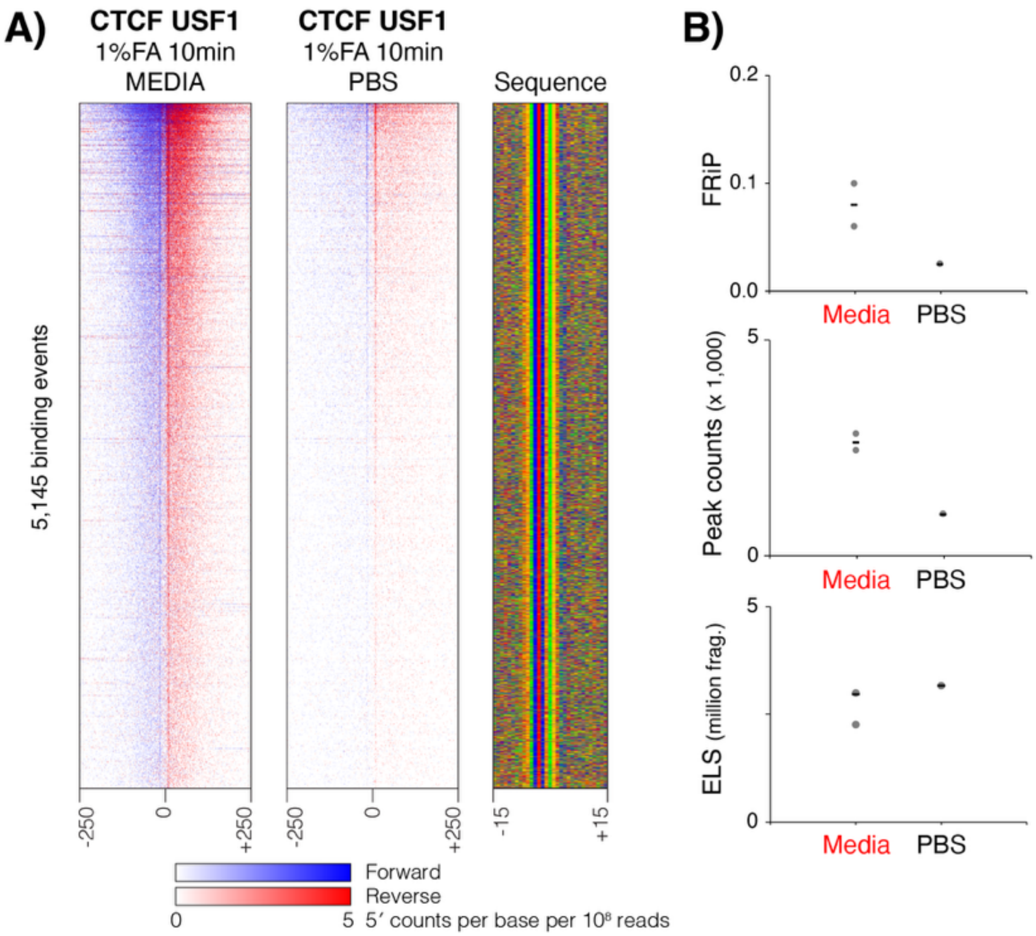


**Supplementary Figure 1. Crosslinking vehicle influences reaction efficiency and USF1 library quality. A)** Heatmaps display normalized per base enrichment of 5′ read positions from ChIP-exo libraries constructed from cells crosslinked in media (left) and PBS (right). The heatmaps plot enrichment in 500bp windows centered on all ChExMix USF1 peaks. Blue represents forward strand 5′ enrichment, while red represents reverse strand 5′ enrichment. **B)** Library quality metrics from USF1 ChIP-exo libraries constructed from cells crosslinked in media and PBS, including FRiP scores, peak counts, and estimated library sizes. Each data point represents an individual ChIP-exo library and short lines represent the mean value across replicates.

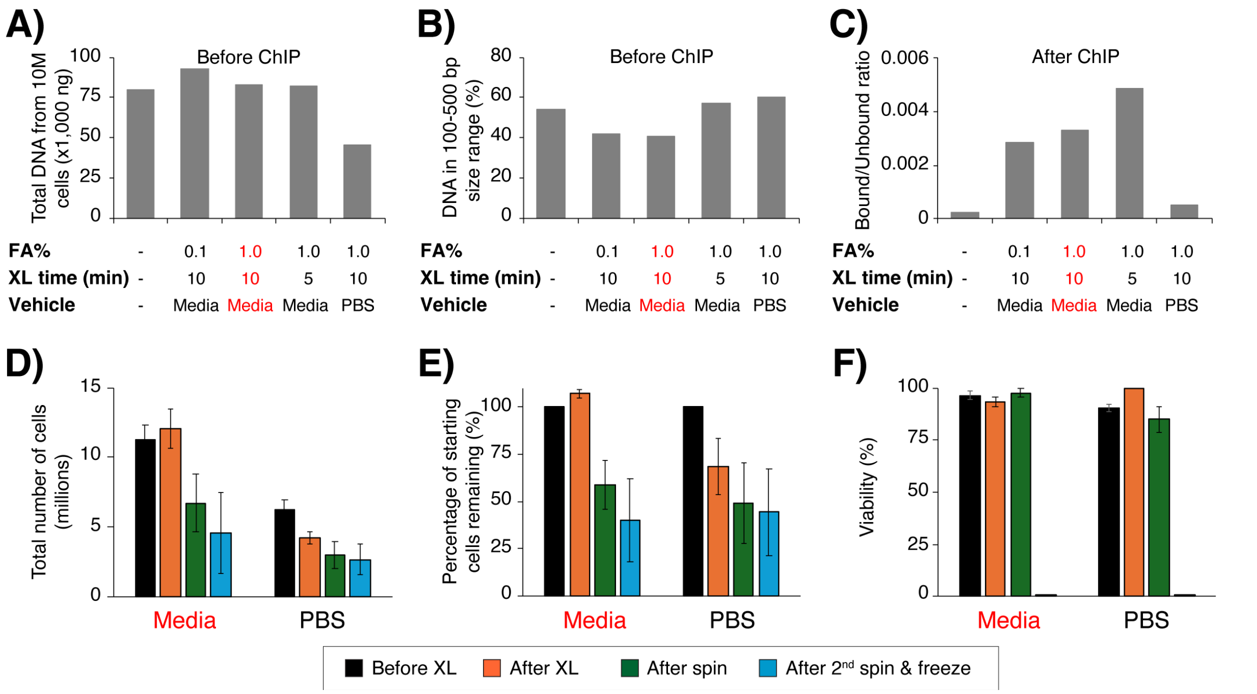


**Supplementary Figure 2. Crosslinking vehicle affects cell yield. A)** Quantification of total DNA before ChIP, as measured by an Agilent Tape Station. **B)** Percentage of total DNA (before ChIP) in the 100-500bp region, as measured by an Agilent Tape Station. **C)** Ratio of bound/unbound DNA after CTCF-ChIP, as measured by a Qubit fluorometer. **D)** Total number of cells per sample during crosslinking to determine yield. **E)** Number of cells per sample, represented as a percentage of total counts. **F)** Percentage cell viability at each step during crosslinking and harvesting. MO-ChIP-exo conditions are highlighted throughout in red.

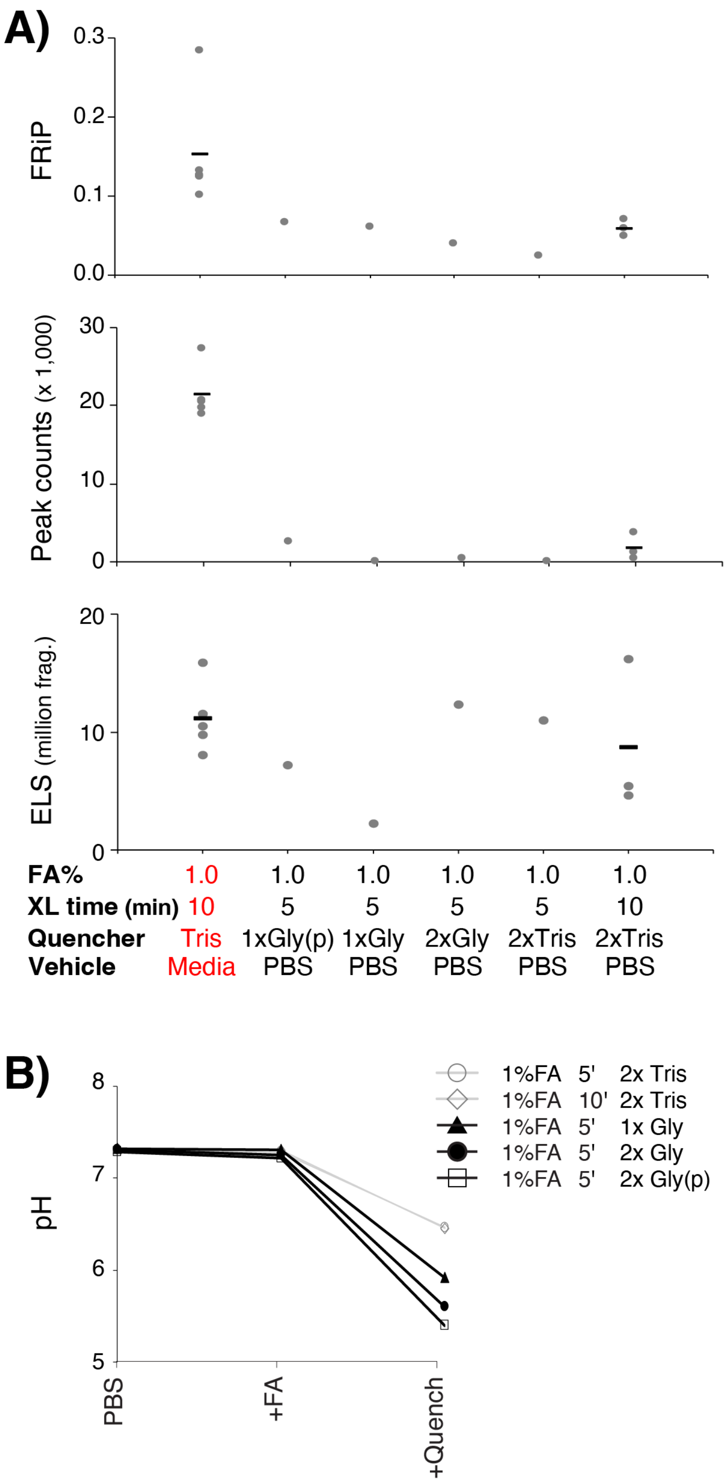


**Supplementary Figure 3. Altering quenching conditions does not mitigate the negative effects of crosslinking in PBS. A)** Quality metrics for CTCF ChIP-exo libraries that were crosslinked in PBS for 5 or 10 min, using Glycine or Tris as quenchers. MO-ChIP-exo conditions are highlighted in red. **B)** pH for samples crosslinked in PBS, before and after crosslinking and quenching, using multiple crosslinking conditions.

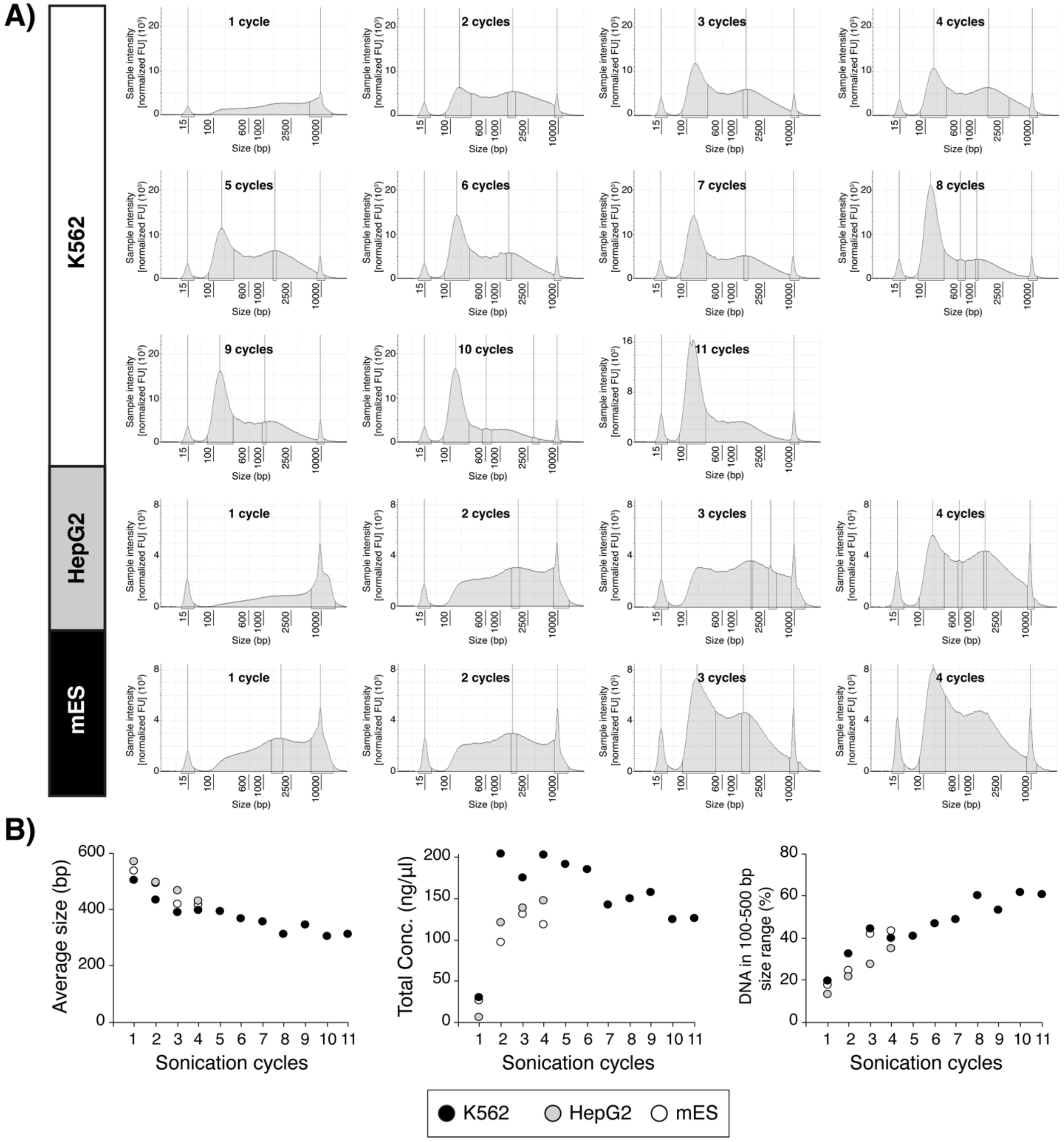


**Supplementary Figure 4. Analysis of DNA fragment size distributions from reverse crosslinked chromatin after sonication. A)** DNA from reverse crosslinked sheared chromatin was analyzed using an Agilent Tape Station across a range of sonication cycles. Electropherograms are shown for K562 samples prepared with between 1 and 11 cycles of sonication, HepG2 samples prepared with between 1 and 4 cycles of sonication, and mESC samples prepared with between 1 and 4 cycles of sonication. **B)** Data from the electropherograms is used to plot the average fragment size, total DNA concentration, and percentage of total DNA in the 100-500 bp region across the relevant titration of sonication cycles.

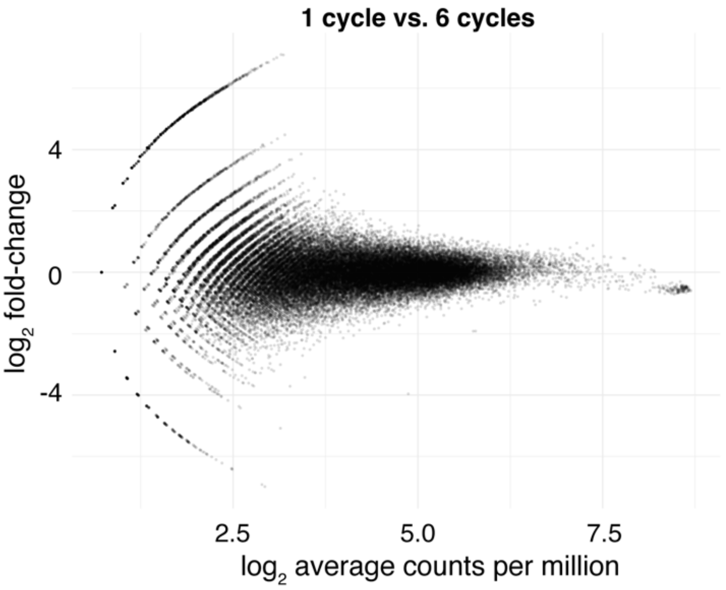


**Supplementary Figure 5. Comparison of CTCF ChIP-exo enrichment levels across sonication cycles.** The MA plot compares enrichment levels between CTCF ChIP-exo libraries prepared with 1 and 6 cycles of sonication. Each dot represents a CTCF binding site. The X axis displays $\log_2$ of the average normalized read counts across conditions at each site. The Y axis displays the $\log_2$ of the fold-change in normalized read counts between the two conditions (i.e., the read count in 1-cycle divided by the read count in 6-cycles).

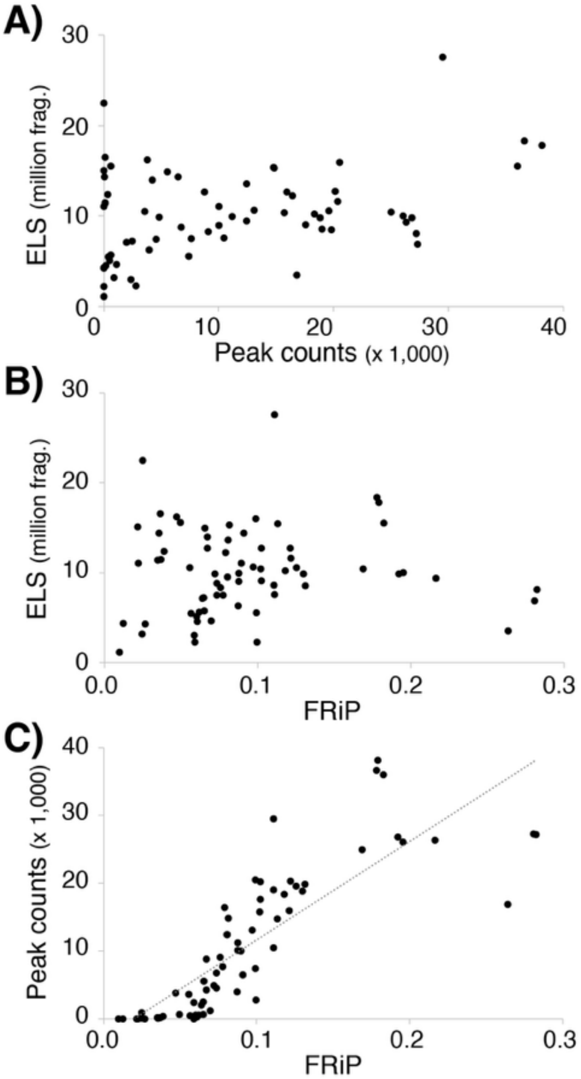


**Supplementary Figure 6. Correlation between the various quality metrics across all libraries included in this study. A)** ELS compared with peak-counts. **B)** ELS compared with FRiP. **C)** Peak counts compared with FRiP. Dots represent individual ChIP-exo experiments in all plots.
