## Supplemental Protocol for "Optimized ChIP-exo for mammalian cells and patterned sequencing flow cells"

### MO-ChIP-exo – Mahony Lab

#### Day 1 - Crosslinking and Harvest

Cells are crosslinked to preserve DNA-protein interactions and harvested, taking precautions to preserve cell and chromatin integrity.

1.1. Count cells (detach first if using adherent cells).
1.2. Adjust cell density to 1M cells/mL with fresh media.
1.3. Choose the size (allow ample space for gentle shaking - wide bottom is preferred) and number of flasks to be used based on the volume. Example: 1 Lt Erlenmeyer for 200 M cells in 200 mL media.

Optional: Crosslinking in an alternative vehicle (i.e., PBS)
*Use caution and count cells directly in crosslinking vehicle without any further centrifugation to ensure an accurate starting number of cells.*

- Centrifuge suspension cells (500 xg for 5 min at RT).
- Aspirate and discard the supernatant.

Note: If using adherent cells, resuspend in vehicle instead of media after detaching cells.

- Resuspend cell pellets in crosslinking vehicle, count cells and adjust cell density to 1M cells/mL.

1.4. Add <u>formaldehyde (FA)</u> to a final concentration of 1%. To prevent oxidation and contamination, the use of ampules is recommended.
*Example: Use 1.5 mL of 32% FA for 50 M cells in 50 mL of media (or vehicle).*
1.5. Incubate with gentle shaking at RT for 5-10 min.
1.6. Quench the crosslinking reaction by adding <u>3 M Tris-HCl pH 7.8</u> in a 2-fold molar excess. Alternatively, use <u>2.5 M Glycine</u> to a final concentration of 125 mM.
*Example: Use 12.5 mL of 3 M Tris-HCl pH 7.8 (or 2.5 mL of 2.5 M Glycine) for 50 mL of crosslinking reaction.*
1.7. Continue to incubate with gentle shaking at RT for 5 min.
1.8. Divide the crosslinked cell suspension into 50 mL tubes and centrifuge at 1,500 xg for 5 min at 4°C in a pre-chilled centrifuge.

*Keep tubes on ice from this point.*

1.9. Discard the supernatant and resuspend the cells in ice-cold PBS + 1X CPI.
1.10. Transfer to microfuge tubes in volumes corresponding to 10-50M cells per tube (or adjust to desired cell number) for freezing.
1.11. Pellet cells by centrifuging for 3min at 10k rpm at 4°C.
1.12. Remove and discard supernatant and immediately flash freeze pellets in liquid nitrogen.
1.13. Store at -80°C until ready to use.

Reagent/consumable List:

- PBS: VWR Cat# VWRL0119-0500
- Protease Inhibitor Cocktail Set V, EDTA-Free, Calbiochem® (CPI): Millipore Sigma Cat# 539139-10VL.
- 32% Formaldehyde: VWR Cat# 100504-858
- Tris: Neta Scientific Cat# RPI-T60040-1000.0
- Glycine: Fisher Scientific Cat# AC120070010

#### Day 2 - Lysis and sonication

Cell and nuclear lysis are performed. The released chromatin is sheared through sonication and checked using a fragment analyzer.

##### Lysis

2.1. Turn on water cooler ~30min before sonicating.
2.2. Thaw crosslinked cell pellets on ice for 10-30min.
2.3. Resuspend 10-30 M cells in 1 mL of <u>CLB + 1X CPI.</u>
2.4. Incubate on a rototorque for 10min at 4°C.
2.5. Needle lyse (25 G needle) for 10 syringe pumps to assist in cell lysis.
2.6. Incubate on a rototorque for 10min at 4°C.
2.7. Spin nuclei at 4°C, 10K rpm, 3 min.
2.8. Remove and discard the supernatant.
2.9. Resuspend the nuclei in 1 mL of <u>NLB + 1X CPI.</u>
2.10. Incubate on a rototorque for 20 min at 4°C.
2.11. Spin at 4°C, 10K rpm, 3 min.
2.12. Remove and discard the supernatant.
2.13. Resuspend chromatin in 300 µL of <u>1X cold PBS + 1X CPI</u> per 10 M cells.
2.14. Needle lyse (25 G needle) for 2 syringe pumps to assist resuspension

##### Sonication

*Note: The first time using a cell line, optimization of shearing conditions is recommended (for the Diagenode Bioruptor® Pico, starting with 1 cycle – with 1 cycle increments – is suggested).*

2.15. Transfer 300 µL of chromatin (from 10M cells) to 1.5 mL Diagenode tubes for sonication.
2.16. Balance the carousels using $dH_2O$ for blanks to complete the spaces if needed.
2.17. Sonicate in a Diagenode Bioruptor® Pico using the 'Go & Shear' setting with the validated parameters on the 'Easy Mode' for 30" ON/OFF for **4 cycles** (or as determined).
2.18. Remove a 15 µL aliquot (~0.5 M cells) to reverse crosslink and purify the DNA for a sonication assessment using the Agilent Tape Station D5000 kit (an alternative fragment analyzer or gel electrophoresis can be used).
2.19. Keep chromatin on ice/cold at all times for short term storage (24-48 hrs) or freeze (-80°C) for long term.

##### Crosslinking Reversal and sonication assessment

2.20. To the 15 µL aliquot removed for sonication assessment, add 185 µL of ChIP Elution Buffer to bring the volume to 200 µL.
2.21. Add 16 µL of 5M NaCl and 1 µL of RNase A (10 mg/mL). Mix well.
2.22. Incubate for 1 hr at 65°C.
2.23. Add 3 µL of proteinase K (20 mg/mL). Mix well.
2.24. Incubate for 1 hr at 45°C.
2.25. Purify the DNA using the Qiagen QIAquick PCR purification kit (or alternative column-based PCR purification kit).
2.26. Assess DNA size distribution using an Agilent TapeStation and D5000 kit. If >30% of DNA is between 100 – 500 bp, proceed to ChIP using the stored chromatin.

Buffer List:

- Cell Lysis Buffer / Modified Farnham Lysis Buffer (CLB) contains 20mM Tris pH 8.0, 85mM KCl, 0.5% NP-40 (Igepal), 0.5% Triton X-100
- Nuclear Lysis Buffer - RIPA Variant (NLB) contains 1x PBS, 1% NP40 (Igepal), 0.5% NaDeoxycholate, 0.1% SDS)
- ChIP Elution Buffer contains 100 mM $NaHCO_3$, 1% SDS
- 5M NaCl

Reagent/consumable List:

- 1.5 mL sonication tubes: Diagenode Cat# No. C30010016
- RNase A (10 mg/mL): Thermo Fisher Scientific Cat# EN0531
- Proteinase K (20 mg/mL): Life Technologies Cat# EO0491
- QIAquick PCR purification kit: Qiagen Cat# 28104
- D5000 ScreenTapes: Agilent Cat# 5067-5588
- D5000 reagents: Agilent Cat# 5067-5589

#### Day 3 - Library Construction: ChIP

Crosslinked sheared chromatin is captured using an antibody of interest to target DNA associated with a specific protein.

*Note: Day 2 and 3 can be combined.*

##### Preparing and Blocking Dynabeads

Note: Determine antibody isotype compatibility with Invitrogen Protein A or G Dynabeads according to the manufacturer's recommendations.

3.1. Keep all buffers, reagents and tubes on ice.
3.2. Mix stock of dynabeads by vortexing and transfer the total volume of beads (50 µL per ChIP) to be used to a 1.7 mL microfuge tube pipetting slowly.
3.3. Place the tube on the magnet for 1 min to collect the beads.
3.4. Remove and discard the supernatant.
3.5. Remove the tube from the magnet and add 1 mL of IPDB + 1X CPI.
3.6. Place the tube on the magnet for 1 min to collect the beads and remove the supernatant.
3.7. Remove the tube from the magnet and add 50 µL of blocking solution (IPDB + 1X CPI + 0.2 mg/mL tRNA).
3.8. Mix and place on a rototorque at 4°C for 15 min.
3.9. Proceed to antibody coating or store at 4°C until ready to use.

##### Coating Dynabeads with specific antibodies and chromatin pre-clearing

3.10. Add 3-10 µg of antibody to the tube containing 50 µL of blocked Protein A/G Dynabead slurry.
3.11. Mix by flicking and incubate for 1-2 hrs on a rototorque at 4°C.

~30min before antibody coating is complete, pre-clear the chromatin to lower the amount of nonspecific binders in the cell lysate:

3.12. If frozen, thaw sonicated crosslinked chromatin for 5 min at 37°C. Otherwise remove from fridge and directly centrifuge at max speed (~14 K) for 15 min at 4°C.

3.13. Once antibody coating is complete, spin tubes in a minifuge to remove any condensation.
3.14. Place tubes against the magnet for 1 min to collect beads.
3.15. Remove and discard the supernatant.
3.16. Immediately add 200 µL of IPDB + 1X CPI.
3.17. Repeat the spin and magnet placement to remove the supernatant.

##### Chromatin Immunoprecipitation (ChIP)

3.18. Immediately add the pre-cleared chromatin to each tube containing antibody/bead complexes. *Optional: save 5 µL from the pre-cleared chromatin to be used as INPUT.*
3.19. Incubate on a rototorque at 4°C **overnight**.

Buffer List:

- IP Dilution Buffer (IPDB) contains 20 mM Tris, pH 8.0, 2 mM EDTA, pH 8.0, 150 mM NaCl, 1% Triton x-100)

Reagent/consumable List:

- Protein A Dynabeads: Invitrogen Cat# 10002D
- Protein G Dynabeads: Invitrogen Cat# 10004D
- tRNA: Millipore Sigma Cat# R8508-1ML

#### Day 4 - Library Construction: First adapter ligation, fill-in, exonuclease digestion, and reverse crosslinking

Antibody-bound chromatin is subjected to end-repair and A-tailing before the first adapter is ligated (using TA ligation). The first adapter is filled in before exonuclease digestion takes place. Chromatin fragments are then reverse crosslinked.

**Washes:**

Before starting the library construction, bead-bound complexes are washed with:

Wash Buffer 1: FA Lysis + 1X CPI
Wash Buffer 2: NaCl 250 + 1X CPI
Wash Buffer 3: LiCl 250 + 1X CPI
Wash Buffer 4: 10 M Tris-HCl pH 8 + 0.4 µL of 25% Triton-X at per mL of buffer

4.1. Spin the tubes on a minifuge and place the tubes on a magnet for 1 min.
4.2. Remove and discard the supernatant (unbound portion).
4.3. Add 150 µL of wash buffer and incubate for 1 min.
4.4. Spin the tubes on a minifuge and place the tubes on a magnet for 1 min.
4.5. Remove wash buffer.
4.6. Repeat for each Wash Buffer (1-4) in the order above.
4.7. After the final wash, directly add the A-tailing master mix.

**A-tailing:**

4.8. Prepare the A-tailing master mix (60 µL per sample) according to the table below:

| Reagent | Vol. per sample (µL) |
|---|---|
| 1X TE | 50 |
| NEBNext® Ultra II End Prep Enzyme Mix | 3 |
| NEBNext® Ultra II End Prep Reaction Buffer | 7 |
| **Total** | **60** |

4.9. Add 60 µL of the A-tailing master mix to the washed beads and gently pipette up/down 10 times.
4.10. Incubate for 30 min at 20°C followed by 30 min at 65°C.
4.11. Proceed directly to ligation.

**First Adapter Ligation (i# adapters):**

4.12. Prepare the First Adapter (i#) Ligation master mix according to the table below:

| Reagent | Vol. per sample (µL) |
|---|---|
| NEBNext® Ultra™ II Ligation Master Mix | 30 |
| NEBNext® Ligation Enhancer | 1 |
| PCR grade $H_2O$ | 1.5 |
| **Total** | **32.5** |

4.13. Quick spin tubes to collect the 60µL at the bottom and directly add 32.5 µL of the First Adapter (i#) Ligation master mix to all tubes.
4.14. Immediately add 1 µL of the first adapter ExB2_i##/ExA2B (15 uM) to each library.
4.15. Gently pipette up/down 10 times to mix and incubate for 15 min at 20°C.
4.16. Wash bead-bound chromatin with Wash Buffers 2-4 as described under ‘Washes’.

**Fill-in Reaction:**

4.17. Prepare the master mix for the Fill-in reaction according to the table below:

| Reagent | Vol. per sample (µL) | Final Concentration |
|---|---|---|
| Water | 24.6 | |
| 10X BSA (1 mg/mL) | 8 | 200 ug/mL |
| 10X Phi29 DNA Polymerase Reaction Buffer | 4 | 1X |
| 3 mM dNTP's | 2.4 | 180 uM |
| Phi29 DNA Polymerase (10 U/µL) | 1 | 10 U |
| **Total** | **40** | |

4.18. Quick spin tubes on a microfuge and place them on a magnet for 1 min.
4.19. Remove and discard the supernatant.
4.20. Add 40 µL of the Fill-in master mix and gently pipette up/down 10 times to mix.
4.21. Incubate for 20 min at 30°C with shaking at 1k rpm.
4.22. Wash dynabeads with Wash Buffer 4 as described under 'Washes'.

**Lambda exonuclease digest:**

4.23. Prepare the Lambda exonuclease master mix according to the table below:

| Reagent | Vol. per sample (µL) | Final Concentration |
|---|---|---|
| Water | 31.6 | |
| 10X Lambda Exonuclease Reaction Buffer | 4 | 1X |
| 10% Triton X | 0.4 | 0.10% |
| DMSO | 2 | 5% |
| Lambda Exonuclease (5 U/µL) | 2 | 10 U |
| **Total** | **40** | |

4.24. Quick spin tubes on a microfuge and place them on a magnet for 1 min.
4.25. Remove and discard the supernatant.
4.26. Add 40 µL of the Lambda exonuclease master mix and gently pipette up/down 10 times to mix.
4.27. Incubate for 30 min at 37°C with shaking at 1k rpm.
4.28. Wash dynabeads with Wash Buffer 4 as described under 'Washes'.

**Elution and Reverse Crosslink:**

4.23 Prepare the ChIP elution Buffer with Proteinase K according to the table below:

| Reagent | Vol. per sample (µL) | Final Concentration |
|---|---|---|
| ChIP Elution Buffer | 40 | |
| Proteinase K (20 mg/mL) | 1.5 | 0.7 mg/mL |

4.24. Quick spin tubes on a microfuge and place them on a magnet for 1 min.
4.25. Remove and discard the supernatant.
4.26. Add 40 µL of ChIP Elution Buffer with Proteinase K and incubate overnight at 65°C.

Buffer List:

- Wash Buffer 1 (FA Lysis Buffer) contains 50 mM HEPES-KOH pH 7.5, 150 mM NaCl, 2 mM EDTA pH 8.0, 0.1% Sodium Deoxycholate and 1% Triton-X 100. Add 1X CPI before use
- Wash Buffer 2 (NaCl 250) contains 50 mM HEPES-KOH pH 7.5, 250 mM NaCl, 2 mM EDTA pH 8.0, 1% Triton-X 100 and 0.1% Sodium Deoxycholate. Add 1X CPI before use
- Wash Buffer 3 (LiCl 250) contains 100 mM Tris-HCl pH 8.0, 250 mM LiCl, 1% NP-40 and 1% Sodium Deoxycholate. Add 1X CPI before use
- Wash Buffer 4 (10 M Tris-HCl pH 8). Add 0.4 µL of 25% Triton-X at per mL of buffer.
- 1X TE contains 10 mM Tris-HCl, pH 8 and 1 mM EDTA
- ChIP Elution Buffer contains 25 mM Tris-HCl pH 7.5, 2 mM EDTA pH 8.0, 200 mM NaCl and 0.5% SDS

Reagent/consumable List:

- NEBNext Ultra II DNA Library Prep Kit for Illumina (Cat# E7645)
- ExB2_i##/ExA2B (15 uM) (Rossi et al. 2018)
- 10X BSA (1 mg/mL) (Thermo Fisher Scientific Cat# J64100-18).
- Phi 29 Polymerase 10 U/µL and 10X Phi29 DNA Polymerase Reaction Buffer (NEB Cat# M0269L)
- dNTP's (Promega Cat# U1330)
- Lambda Exonuclease 5 U/µL and 10 X Lambda Exonuclease Reaction Buffer (NEB Cat# M0262S)
- 10% Triton X (Sigma-Aldrich Cat# X100-1L)
- DMSO (Sigma-Aldrich Cat# D2650-100ML)

#### Day 5 - Library Construction: Second adapter ligation, enrichment and quantification.

DNA fragments undergo 2nd adapter ligation. Libraries are then enriched and size selected before final quantification.

**DNA Purification:**

5.1. Quick spin tubes on a microfuge and place them on a magnet for 1 min.
5.2. Transfer and *KEEP the supernatant* (eluate) in a new tube.
5.3. Add *1.8 Volumes of TotalPure Beads* (72 µL for 40 µL eluted samples).
5.4. Mix by pipetting up and down 10 times and incubate at RT for 5 min.
5.5. Place tubes on a magnet for 1 min and remove the supernatant.
5.6. Keep tubes on magnet and add 180 µL of freshly made 70% ethanol without disturbing the pellet.
5.7. Incubate for 30 sec without disturbing the beads.
5.8. Remove the supernatant and repeat the ethanol wash.
5.9. Dry the beads at RT for 5 min.
5.10. Add 21 µL of PCR-grade $H_2O$ and mix by pipetting up and down 10 times.
5.11. Place tubes on a magnet and collect 20 µL of the eluted sample in a fresh tube.

**Second Adapter Ligation (iX adapters):**

5.12. Prepare the Second Adapter (iX) Ligation master mix according to the table below:

| Reagent | Vol. per sample (µL) | Final Concentration |
|---|---|---|
| Water | 9 | |
| 10X T4 Ligase Buffer | 4 | 1X |
| T4 Ligase (600 U/µL) | 2 | 1200 U |
| **Total** | **15** | |

5.13. Add 15 µL of the Second Adapter (iX) Ligation master mix to each tube containing 20 µL of eluted DNA.
5.14. Immediately add 5 µL of the second adapter ExB1-iX/ExB1B-5N (3 uM) to each library.
5.15. Gently pipette up/down 10 times to mix and incubate for 1 hr at 25°C.

**Repeat DNA Purification:**

5.16. Repeat steps 5.1 – 5.11 to clean up DNA fragments containing both adapters

**PCR:**

5.17. Prepare the PCR Master Mix (plan to include negative and positive controls) according to the table below:

| Reagent | Vol. per sample (µL) | Final Concentration |
|---|---|---|
| Water | 6.8 | |
| 5X Phusion HF Buffer | 8 | 1X |
| 3 mM dNTP's | 2.67 | 200 uM each |
| P1.3 primer (20 uM) | 0.8 | 500 nM |
| P2.1 primer (20 uM) | 0.8 | 500 nM |
| Phusion Hot Start DNA Polymerase (2 U/µL) | 1 | 2 U |
| **Total** | **20** | |

5.18. Add the 20 µL of master mix to each library (20 µL resuspended DNA).
5.19. Vortex tubes and spin down before placing them in the thermocycler using the following settings:

Thermocycler settings

| | | | |
|---|---|---|---|
| Denature/Phusion Activation | | 98°C | 1 min |
| Amplification (12 cycles) | Denature | 98°C | 10 sec |
| | Anneal | 52°C | 30 sec |
| | Extension | 72°C | 30 sec |
| End Extension | | 72°C | 2 min |
| | | 4°C | Hold |

**Double-sided Size selection:**
This step uses bead to sample ratios of 0.7X and 1X to select for ~150 - 500 bp fragments
An alternative to this step is gel excision and purification as in ChIP-exo 5.0

5.20. Spin tubes to collect libraries at the bottom of the tube.
5.21. Add 10 µL 10 mM pH 7.5 to each 40 µL library for a total starting volume of 50 µL.
5.22. Gently resuspend SPRI beads.
5.23. Add 0.7 volumes (35 µL) of beads (50 µL * 0.7x vol) to each library.
5.24. Mix by pipetting 10 times and incubate at RT for 5 min.
5.25. Place samples on magnet and allow the beads to settle for 1-3 min.
5.26. Collect the supernatant (contains the Right Side Size Selected sample) and transfer to fresh tubes.
5.27. Gently resuspend fresh SPRI beads.
5.28. Add 0.3 volumes (15 µL) of beads. Calculated as: Sample Vol. * (1X – initial ratio (0.7X)). [50 µL * (1-0.7) = 15 µL of SPRIselect beads]
5.29. Mix by pipetting 10 times and incubate at RT for 5 min.
5.30. Place samples on magnet and allow the beads to settle for 1-3 min.
5.31. Remove and discard the supernatant.
5.32. With the tubes still on the magnet, add 180 µL of freshly made 85% ethanol and incubate at RT for 30 sec.
5.33. Remove and discard the supernatant. Be careful not to remove any beads at this step as it may result in reduced yield.
5.34. Repeat the ethanol wash.
5.35. Air dry pellet for 5 min.
5.36. Remove the samples from the magnet and resuspend the pellet in 21 µL of 10 mM Tris pH 8.
5.37. Mix by pipetting 10 times and incubate for 5 min.
5.38. Place samples back on the magnet and allow the beads to settle for 1-3 min.
5.39. Transfer 20 µL of the supernatant to fresh tubes.

**DNA Purification:**
5.40. Repeat steps 5.1 – 5.11 to clean up libraries and remove any excess adapters or adapter dimers remaining.

**Library assessment:**
5.41. Assess DNA size distribution using an Agilent Tape Station and High Sensitivity D5000 kit. >90% of each library is expected to be between 200-500 bp with an average fragment size of 300-350 bp. It's important to also confirm the absence of free adapters (~75 bp) or adapter dimers (~150 bp).

5.42. Finally, library quantification is performed using NEB Next® Library Quantification Kit, For Illumina® in an ABI StepOne Plus Real Time PCR System.

Reagent/consumable List:

- Mag-Bind® TotalPure NGS (Omega Bio-Tek M1378-01)
- T4 DNA Ligase MBG 500 U and 10x T4 Ligation Buffer (Qiagen Cat no. / ID. EN11-050)
- ExB1-iX/ExB1B-5N (3 uM) from Rossi et al. 2018
- Phusion™ High-Fidelity DNA Polymerase (2 U/µL ) and 5X HF Buffer (Thermo Fisher Scientific Cat# F549L)
- PCR primers 1.3 and 2.1 from Rossi et al. 2018
- SPRIselect DNA Size Selection reagent (Beckman Coulter B23317)
- HS D5000 ScreenTapes (Agilent Cat# 5067-5592)
- HS D5000 reagents (Agilent Cat# 5067-5593)
- NEB Next® Library Quantification Kit, For Illumina® (NEB Cat# E7645L)

### Additional Details

**Equipment/Materials required:**

- Rototorque at 4°C
- Minifuge
- Vortex
- Benchtop centrifuge
- Refrigerated benchtop centrifuge
- Magnet for collection of dynabeads
- Heatblock and thermomixer
- Thermocycler
- Agilent 2100 Bioanalyzer Instrument or equivalent fragment analyzer
- ABI StepOne Plus Real Time PCR System (or similar)

**Antibodies:**

The following antibodies were used successfully for MO-ChIP-exo:

| Antibody | Catalog Number |
|---|---|
| Anti-CTCF | EMD Milipore 07-729 |
| IgG | Sigma i5006 |
| Anti-USF1 | DSHB USF1-1B8 |

**Adapter pre-annealing:**

ChIP-exo oligos are single stranded and need to be pre-annealed for each adapter as below. For MO-ChIP-exo oligo's are HPLC purified and pre-annealed adapters are used as UDI's.

ExB2-iX (Number Adapter i01– i16) – 15 uM

- Mix ExA2B with each ExB2-i# (i01– i16) (15 uM each) in 1X SSC.
- Incubate 2 min at 95°C followed by 1 hour at room temp.
- Store at -20C until ready to use.
-

ExC1-iX (Letter Adapters iA-iP) – 3 uM

- Mix ExA1-SSL_N5 with each ExC1-iX (iA-iP) (15 uM each) in 1X SSC.
- Incubate 2 min at 95°C followed by 1 hour at room temp.
- Dilute 1:5 in 10 mM pH 7.5 for 3 uM working stocks.
- Store at -20C until ready to use.
